# *Bacteroides thetaiotaomicron* uses a widespread extracellular DNase to promote bile-dependent biofilm formation

**DOI:** 10.1101/2021.06.04.447082

**Authors:** Nathalie Bechon, Jovana Mihajlovic, Anne-Aurélie Lopes, Sol Vendrell-Fernández, Julien Deschamps, Romain Briandet, Odile Sismeiro, Isabelle Martin-Verstraete, Bruno Dupuy, Jean-Marc Ghigo

## Abstract

*Bacteroides thetaiotaomicron* is a gut symbiont that inhabits the mucus layer and adheres to and metabolizes food particles, contributing to gut physiology and maturation. Whereas adhesion and biofilm formation could be key features for *B. thetaiotaomicron* stress resistance and gut colonization, little is known about the determinants of *B. thetaiotaomicron* biofilm formation. We previously showed that the *B. thetaiotaomicron* reference strain VPI-5482 is a poor *in vitro* biofilm former. Here we demonstrated that bile, a gut-relevant environmental cue, triggers the formation of biofilm in many *B. thetaiotaomicron* isolates and common gut Bacteroidales species. We identified the genetic determinants of this bile-dependent biofilm formation and showed that it involves the production of the DNase BT3563, degrading extracellular DNA, in biofilms formed in the presence of bile. Our study therefore identifies a physiologically relevant condition inducing *B. thetaiotaomicron* biofilm and shows that, in contrast to the biofilm-promoting role played by bacterial eDNA scaffold in Firmicutes and Proteobacteria models, degradation of eDNA by BT3563 DNAse and its widespread homologs is required to achieve *B. thetaiotaomicron* bile-dependent biofilm formation.

## INTRODUCTION

*Bacteroides thetaiotaomicron*, one of the most abundant bacterial gut symbionts, is involved in the degradation of complex polysaccharides and the maturation of the host immune system (Eckburg et al., 2005; Martens et al., 2008; Wexler, 2007). *B. thetaiotaomicron* colonizes the gel-like mucus layer of intestinal epithelia cells (IECs) in both healthy and diseased conditions, and also adheres to food particles (Macfarlane and Macfarlane, 2006; Mark Welch et al., 2017; Sonnenburg et al., 2005), suggesting that adhesion to host surfaces and the formation of bacterial biofilm aggregates could play an important biological role. Consistently, the transcriptional profile of *B. thetaiotaomicron* colonizing the mouse gut is more similar to *B. thetaiotaomicron in vitro* biofilm than to *in vitro* planktonic culture (TerAvest et al., 2014). Whereas the reference strain VPI-5482 only forms poor biofilms *in vitro*, several mutations leading to increased aggregation and biofilm capacity *in vitro* were recently identified, revealing the importance of capsular polysaccharides and type V pilus in mediating adhesion (Béchon et al., 2020a; Mihajlovic et al., 2019). Moreover, polysaccharide utilization loci (PUL) have been proposed to mediate *Bacteroides* adhesion to substrates. PULs are surface structures involved in sugar degradation that are composed of at least one TonB-dependent receptor and one cell surface lipoprotein binding a glycan substrate (Grondin et al., 2017; Martens et al., 2008). Mucin-O-glycan degrading PULs might mediate adhesion to mucin and therefore favor mucus colonization in the gut upon bile stimulation (Martens et al., 2008; TerAvest et al., 2014). Although several gut environmental cues and compounds including antibiotics, oxygen, metal concentration, mucins, immunoglobulin A and bile (Ahn and Burne, 2007; Bollinger et al., 2001, 2006; Dubois et al., 2019; Hoffman et al., 2005; Pumbwe et al., 2007; Sicard et al., 2017) were shown to modulate bacterial biofilm capacity in various bacterial species, gut-relevant conditions promoting *B. thetaiotaomicron* biofilm formation are still poorly understood.

In the closely related *Bacteroides fragilis* species, both sub-inhibitory concentrations of the antibiotic enrofloxacin and bile were independently shown to increase *in vitro* biofilm formation, with consequences on *B. fragilis* physiology (Pumbwe et al., 2007; Silva et al., 2014). In this study, we showed that exposure to sub-inhibitory concentrations of bile extract induced biofilm formation in *B. thetaiotaomicron* strain VPI-5482 and in the majority of tested *B. thetaiotaomicron* clinical isolates and several common gut Bacteroidales species. Using transposon mutagenesis, we identified a *B. thetaiotaomicron* VPI-5482 operon, *BT3560-3563,* involved in bile-dependent biofilm formation. We showed that BT3563 is an extracellular DNase that is well-represented in *B. thetaiotaomicron* species and that it degrades extracellular DNA (eDNA) in biofilms formed in the presence of bile. Our study therefore shows that, in contrast to the biofilm-promoting role of bacterial extracellular DNA in many species (Okshevsky and Meyer, 2015), degradation of eDNA by BT3563 or by other extracellular nucleases may be a widespread requirement to achieve biofilm formation in presence of bile, as shown with *B. thetaiotaomicron*.

## RESULTS

### Bile induces biofilm formation in *B. thetaiotaomicron* and several Bacteroidales

To identify gut-relevant conditions inducing biofilm formation in *B. thetaiotaomicron* VPI-5482, we supplemented BHIS growth medium with various components of the normal intestinal environment, including sugars (0.5% D-glucose, D-mannose, D-rhamnose, D-cellobiose and D-maltose), hemin (25 mg/L and 50 mg/L), mucin (0.1% and 0.5%) and a mix of bovine - ovine bile extract (0.1%, 0.5%, 1%) (hereafter referred to as bile) for 24h (Supplementary Figure S1). We showed that addition of bile increased biofilm formation at 24h without impacting viable cell count (minimal inhibitory concentration for bile is 4%) (Supplementary Figure S1 and S2). This increase was maximal at 48h in the presence of 0.3 to 1% bile (Figure 1A). We then tested biofilm formation at 48h in the presence of 0.5% bile in a collection of *B. thetaiotaomicron* clinical isolates (Mihajlovic et al., 2019) (Supplementary Table S1) and a representative strain of the most common gut *Bacteroides* and *Parabacteroides* species. We showed that, except for *B. thetaiotaomicron* strain jmh71 and the tested *B. vulgatus* strain, all strains displayed an increased biofilm phenotype, indicating that bile is a widespread inducer of biofilm formation amongst gut Bacteroidales species (Figure 1BC).

**Figure 1.**
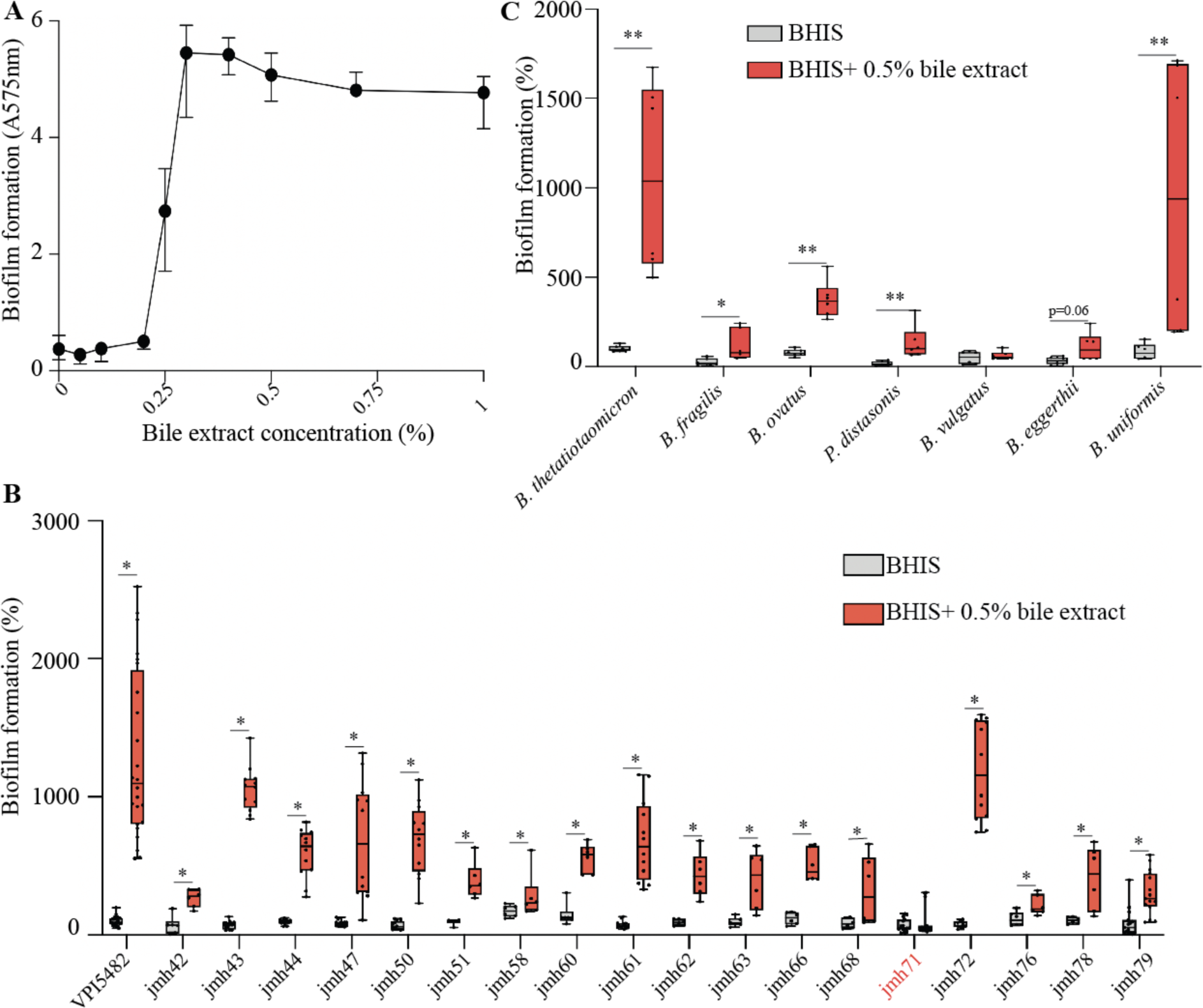
Bile extract promotes biofilm formation of several Bacteroidales isolates. **A.** *B. thetaiotaomicron* VPI-5482 48h biofilm formation as a function of bile extract concentration. Mean of 6 biological replicates, error bars represent standard deviation. **B.** and **C.** 96-well plate crystal violet 48h biofilm assay. Mean of *B. thetaiotaomicron* VPI-5482 grown in BHIS is adjusted to 100 %. Min-max boxplot of 6-12 biological replicates for each strain. *, p-value<0.05, ** p-value<0.005, Mann-Whitney test.

### Exposure to bile induced a major transcriptional shift but did not reveal biofilm functions

To understand the impact of bile on *B. thetaiotaomicron* physiology, we performed a RNAseq analysis comparing *B. thetaiotaomicron* grown overnight in presence or absence of 0.5% bile. We found that bile induces a widespread transcriptional shift, with 17% genes showing a significant change (Supplementary Table S2A). In the presence of bile, out of 4816 genes, 378 and 429 genes were significantly up- and down-regulated, respectively (Supplementary Table S2BC). About half of these genes could be assigned a functional COG category, spanning almost all COG categories (Supplementary Table S2D). The 5 genes showing the highest fold change (>50) are organized in two operons, *BT2793-2795*, encoding a bile-specific tripartite multidrug efflux system (Liu et al., 2019), and *BT0691-0692*, predicted to encode an outer membrane protein and a calcineurin superfamily phosphohydrolase, respectively. Consistently, both of these loci were previously shown to be important for fitness under bile salt stress (Liu et al., 2019), and are probably involved in bile tolerance. Although we did not observe any induction of genes encoding capsular polysaccharides (Coyne and Comstock, 2008; Xu et al., 2003) or type V pilin homologs (Xu et al., 2016) that were previously shown to contribute to *B. thetaiotaomicron* biofilm formation (Béchon et al., 2020b; Mihajlovic et al., 2019), several polysaccharide utilization loci (PUL) involved in mucin O-glycan and host-derived sugars degradation (Martens et al., 2008) that could contribute to adhesion to the mucus layer were up-regulated, suggesting that bile exposure might increase *B. thetaiotaomicron* adhesion capacity also *in vivo*. These results showed that exposure to bile leads to important physiological adaptations in *B. thetaiotaomicron* that might increase its colonization capacity. This transcriptomic approach did not, however, reveal any regulation of genes encoding known *B. thetaiotaomicron* determinants of biofilm formation *in vitro*.

### Identification of an operon involved in bile-dependent biofilm formation

To identify the genetic determinants involved in bile-dependent biofilm formation, we performed a random transposon mutagenesis in *B. thetaiotaomicron* VPI-5482 and screened for mutants failing to produce biofilm in the presence of 0.5% bile. We identified 9 biofilm-negative transposon mutants out of 6510 mutants screened (Figure 2A), all corresponding to transposon insertions in a potential operon (Supplementary Figure S3) encoding a TonB-dependent receptor (*BT3560* - 7 insertions), a hypothetical protein (*BT3561*) and two putative membrane-associated lipoproteins with nuclease activity (*BT3562 -*2 insertions and *BT3563)* (Figure 2B). Whereas deletion of *BT3561* had no impact on biofilm formation, the deletion of *BT3560, BT3562*, *BT3563* or of the whole *BT3560-3563* region significantly reduced bile-dependent biofilm formation compared to WT without associated growth defect (Figure 2C and Supplementary Figure S4). Complementation of *ΔBT3563* and *ΔBT3560-3563* mutants by p*BT3563,* a plasmid constitutively expressing *BT3563*, restored bile-dependent biofilm formation (Figure 2D), whereas *ΔBT3560* or *ΔBT3562* strains were not complemented upon addition of p*BT3560* or p*BT3562*, suggesting that deletion of *BT3560* and *BT3562* might have a polar effect on the downstream *BT3563* gene (Figure 2D). However, since a *ΔBT3560-3563* mutant formed slightly but reproducibly less biofilm than a *ΔBT3563* simple mutant, *BT3560* and *BT3562* might also contribute to biofilm formation in the presence of bile. Taken together, these results showed that *BT3563* is necessary for proper bile-dependent biofilm formation of *B. thetaiotaomicron* VPI-5482.

**Figure 2.**
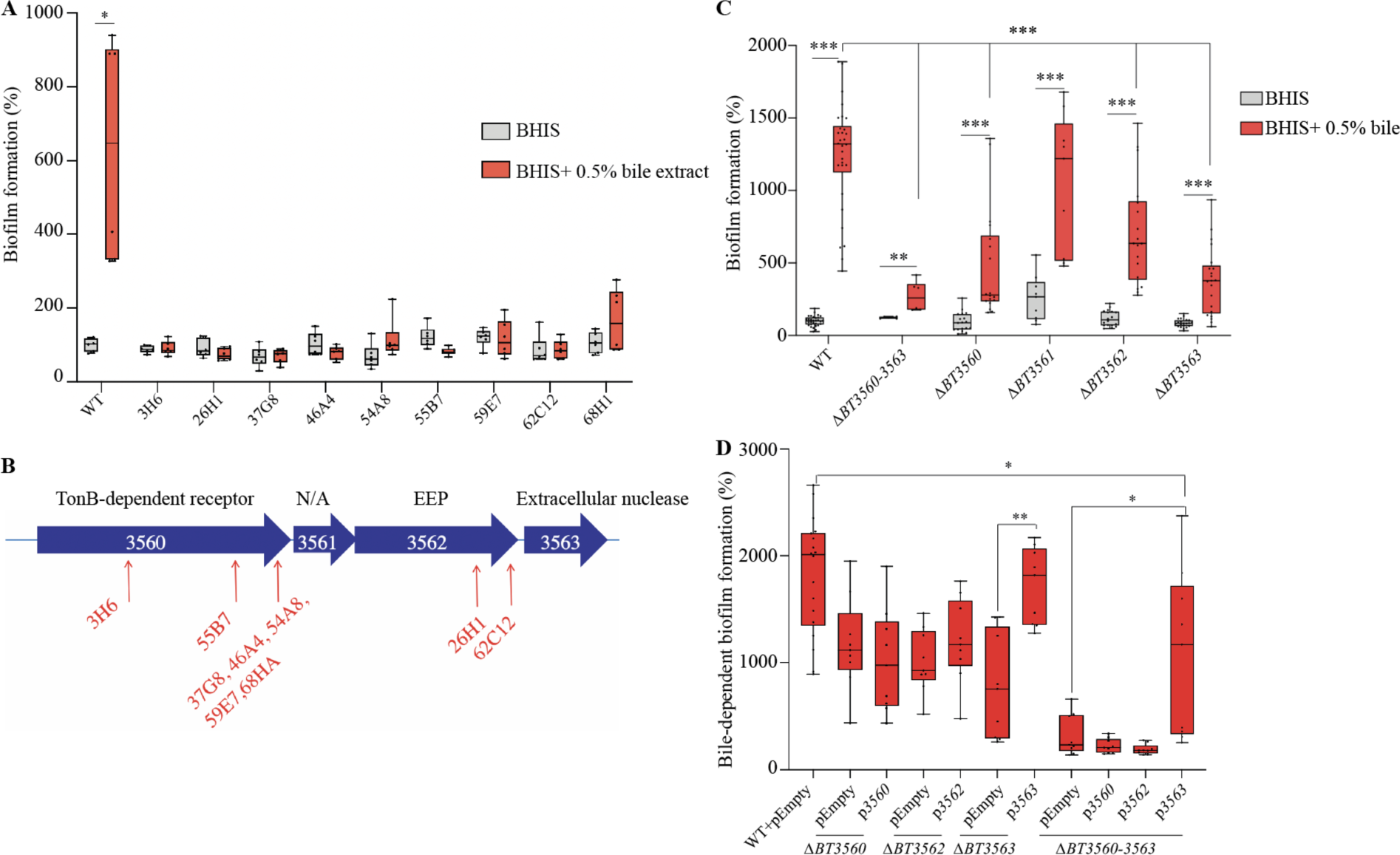
*BT3560-3563* is required for bile-dependent biofilm formation. **A.** and **C.** 96-well plate crystal violet biofilm assay after 48h growth in BHIS or BHIS+0.5% bile extract. **B.** Schema of the *BT3560-3563* genetic locus. Red arrows indicate transposon insertion points. N/A: not annotated, EEP: exonuclease/endonuclease/phosphatase family **D.** 96-well plate crystal violet biofilm assay after 48h growth in BHIS+0.5% bile extract. WT in BHIS is not represented. **A.**, **C.** and **D.** Mean of WT in BHIS is adjusted to 100 %. Min-max boxplot of 6-18 biological replicates for each strain, each replicate is the mean of two technical replicates. * p-value<0.05, ** p-value<0.005, ***p-value<0.0005, Mann-Whitney test.

### *BT3563* is an extracellular DNase degrading eDNA in biofilms formed in presence of bile

To investigate the potential nuclease activity of BT3563, we exposed genomic DNA to *B. thetaiotaomicron* culture supernatant. We showed that DNA was degraded after an overnight incubation with a WT supernatant. This degradation was abolished when we used a *ΔBT3563* strain carrying the empty vector (Figure 3A). Complementation with p*BT3563* plasmid, constitutively expressing *BT3563,* restored DNA degradation, confirming the DNAse activity of BT3563 (Figure 3A). To confirm the contribution of BT3563 extracellular DNase to bile-dependent biofilm formation, we complemented the absence of BT3563 in *ΔBT3560-3563* and *ΔBT3563* strains by adding purified DNase I to the growth medium. We observed an increase of bile-dependent biofilm formation in *ΔBT3560-3563* and *ΔBT3563* strains in presence of DNase I, but a decrease in WT biofilm formation (Figure 3B). This prompted us to measure the concentration of extracellular DNA (eDNA) in the extracellular matrix (ECM) of *B. thetaiotaomicron* biofilms formed in the presence of bile. We showed that, whereas deletion of *BT3563* did not significantly impact ECM protein concentration (Supplementary Figure S5), it increased eDNA concentration compared to WT. Upon addition of p*BT3563* in trans, the eDNA concentration was reduced back to WT level, which is consistent with BT3563-dependent degradation of DNA in the ECM (Figure 3C). Addition of bile was not necessary to observe BT3563-mediated DNA degradation (Figure 3A), and our transcriptomic analysis showed no increase of *BT3563* transcription in the presence of bile (Supplementary Table S2A), suggesting that bile might not directly impact BT3563 production or activity. We then assessed the impact of bile on eDNA concentration itself and observed an increase of eDNA concentration in *B. thetaiotaomicron* supernatant of cultures grown in presence of bile, indicating that bile could increase eDNA release (Supplementary Figure S6). These results showed that BT3563 is an extracellular DNase that degrades eDNA in the biofilm matrix and that bile exposure increases the release of eDNA, which might be one of the mechanisms by which bile exposure favors biofilm formation.

**Figure 3.**
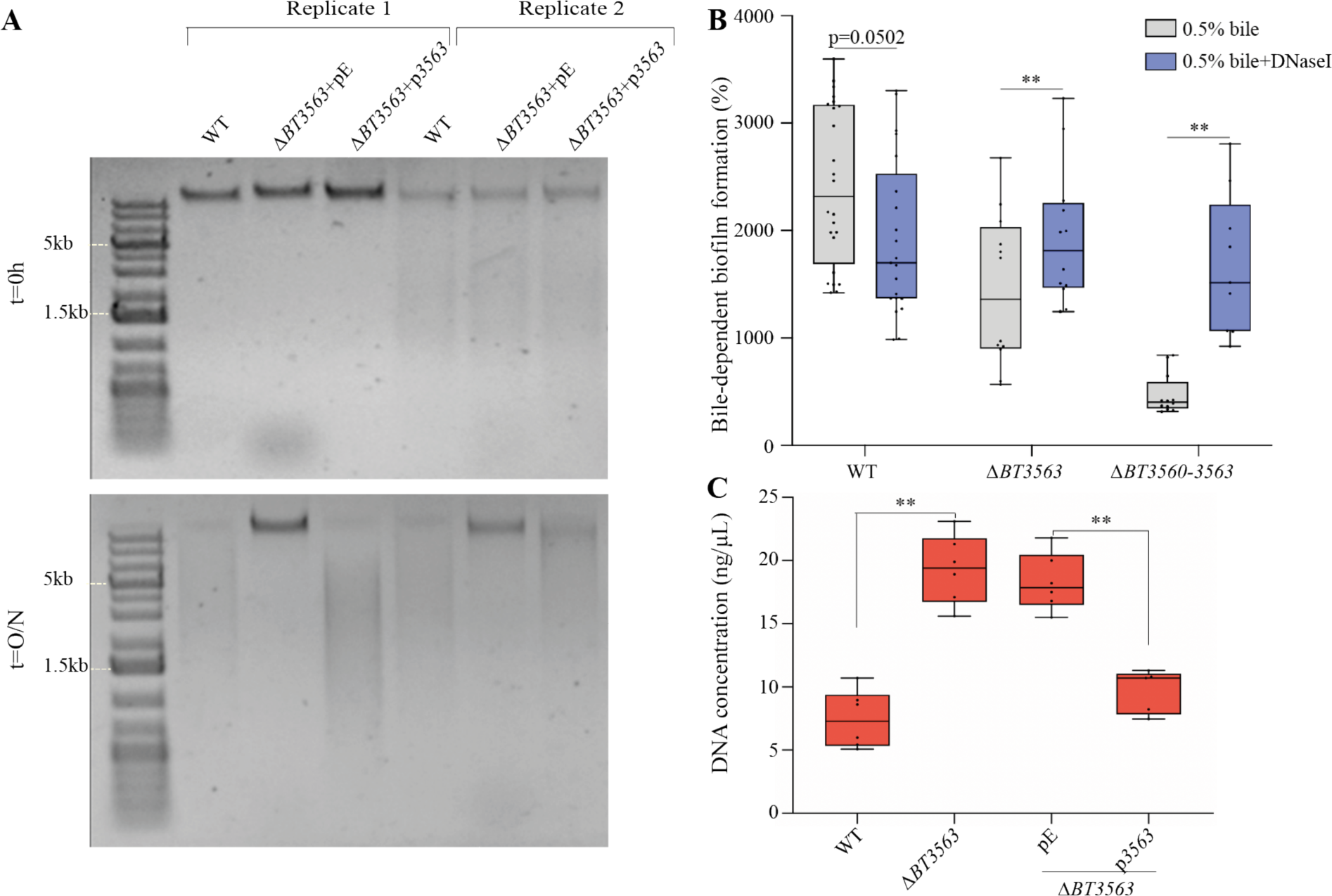
BT3563 degrades nucleic acids in the extracellular matrix of bile-dependent biofilms. **A.** 0.8% agarose gel of the supernatant of overnight cultures grown without bile and mixed with genomic DNA (23ng/µL final concentration) initially (t=0h) and after an overnight incubation at 37°C (t=18h). For each strain, 2 biological replicates are shown. **B.** 96-well plate crystal violet biofilm assay after 48h growth in BHIS+0.5% bile extract, with or without DNase I. Mean of WT in BHIS is adjusted to 100 % (not represented). Min-max boxplot of 9-21 biological replicates for each strain, each replicate is the mean of two technical replicates. * p-value<0.05, *** p-value<0.0005 Wilcoxon matched-pairs signed rank test. **C.** Extracellular DNA concentration (ng/µL) in the purified extracellular matrix of bile-dependent biofilms grown in BHIS+0.5% bile extract for 48h. Min-max boxplot of 6 biological replicates for each strain. * p-value<0.05, ** p-value<0.005, Mann-Whitney test.

### Extracellular DNase activity is necessary to observe maximum bile-dependent biofilm formation

To analyze the distribution and abundance of eDNA in the ECM of *B. thetaiotaomicron* bile-dependent biofilms, we labelled eDNA with TOTO-1 dye and imaged 48h 96-well plate biofilms by confocal laser scanning microscopy (CLSM). For each condition, a representative well is shown in Figure 4, and a picture of the two other wells is shown in Supplementary Figure S7. In presence of bile, *B. thetaiotaomicron* biofilm was expectedly denser and more structured than in absence of bile (Figure 4AB and Supplementary Figure S7). Nucleic acids were more abundant in presence of bile (Supplementary Figure S8) and they were distributed diffusely in the entire biofilm, with spots potentially corresponding to dead cells (Figure 4AB and Supplementary Figure S7). To determine the nature of these nucleic acids we grew *B. thetaiotaomicron* biofilms in presence of bile and DNase I or RNase I. Whereas RNase I treatment had little impact on *B. thetaiotaomicron* biofilms (Figure 4C and Supplementary Figure S7-S8), addition of DNase I strongly decreased the amount of nucleic acids detected (Figure 4D and Supplementary Figure S8), confirming that DNA was the most abundant nucleic acid in the ECM. Addition of DNase I also led to a loss of *B. thetaiotaomicron* biofilm 3D structure, suggesting eDNA might be an important structuring component of the ECM (Figure 4D and Supplementary Figure S7). We then showed that, whereas the *ΔBT3563* mutant formed scattered biofilms in the presence of bile, with dispersed or small clumps of cells (Figure 4EF and Supplementary Figure S7), addition of DNase I during growth of *ΔBT3563* restored a WT homogenous and dense biofilm phenotype (Figure 4DG and Supplementary Figure S7). One hypothesis would be that DNase activity might be required to prevent repulsion between clumps of cells, or between cells and some components of the extracellular matrix, allowing the formation of denser structures. By contrast to eDNA quantification in the extracted ECM, there was no clear increase in TOTO-1 labelling of *ΔBT3563* or *ΔBT3560-3563* mutants compared to WT in the presence of bile (Supplementary Figure S8), which could be due to a lack of sensitivity of eDNA quantification by CLSM. These results demonstrated that DNA, rather than RNA, contributes to biofilm structure in the presence of bile and that the presence of either BT3563 or the addition in culture of DNase I lead to the formation of denser bacterial structures, allowing maximum biofilm formation.

**Figure 4.**
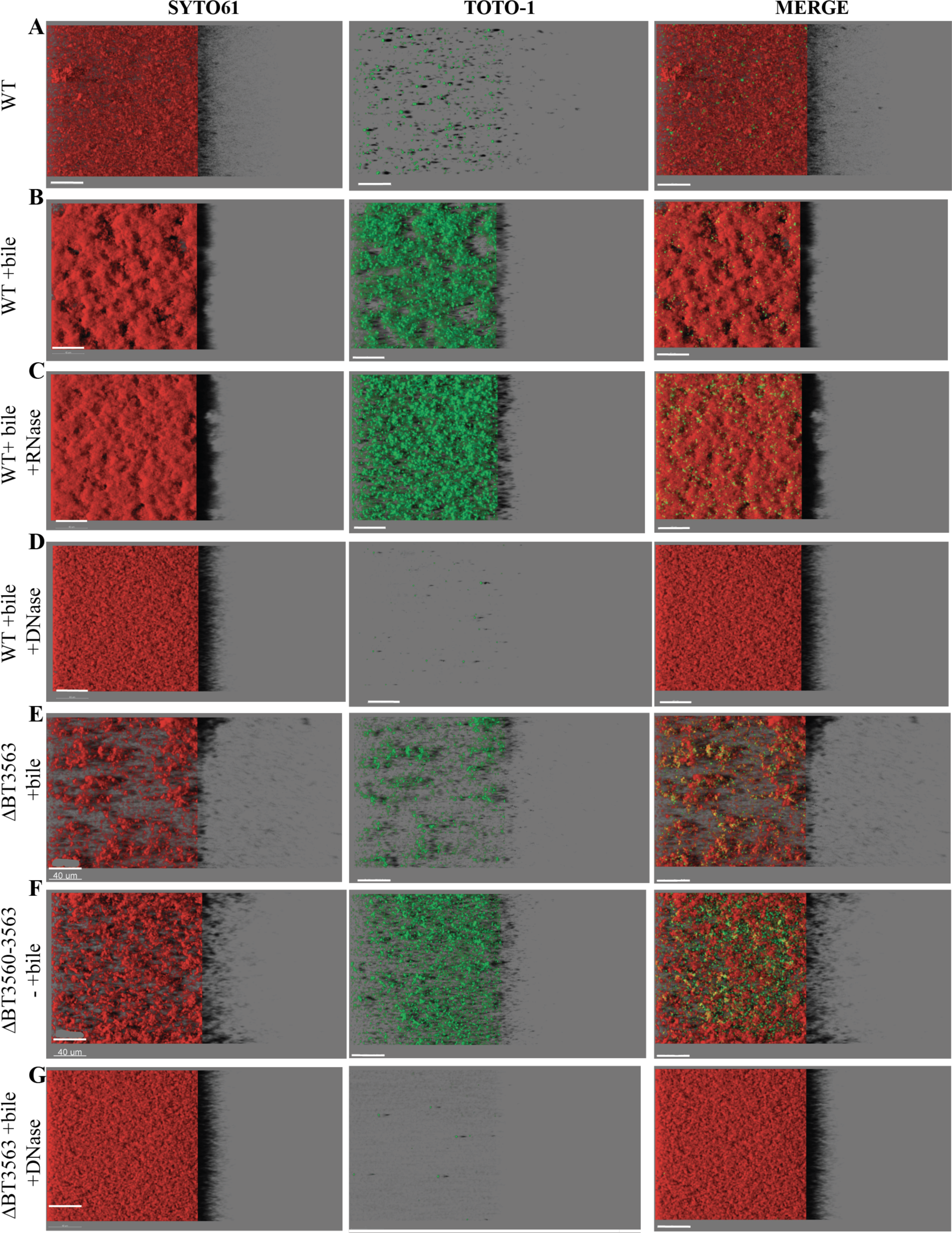
Visualization of *B. thetaiotaomicron* biofilms. IMARIS easy 3D projections from confocal laser scanning (CLSM) microscopy images of *B. thetaiotaomicron* biofilms grown in plates in absence or presence of 0.5% bile extract (“+bile”), RNase I (“+RNase”), or DNase I (“+DNase”). Cells are labelled with SYTO61 dye (left panel, in red). eDNA and dead cells are labelled with TOTO-1 (middle panel, in green). The merged image is shown on the right. For each image, the virtual shadow projection of the biofilm is shown on the right. Scale bars represent 40 µm.

### *BT3563* homologs are involved in bile-dependent biofilm formation of *B. thetaiotaomicron* clinical isolates

To further investigate the correlation between BT3563 and bile-dependent biofilm, we sequenced the whole genome of the 18 *B. thetaiotaomicron* jmh isolates. We found a *BT3563* homolog in all tested strains, even in *B. thetaiotaomicron* jmh71, the only strain that did not form biofilms in the presence of bile (Supplementary Table S3), suggesting additional factors might be responsible for lack of bile-dependent biofilm formation in this strain. In 11 out of the 18 jmh strains (jmh42, 44, 47, 50, 51, 58, 61, 66, 76, 78 and 79), the homology with VPI-5482 BT3563 exceeded 80% homology (Supplementary Table S3) and the genomic region around the detected *BT3563* homolog gene was similar to that of VPI-5482 and included homologs of genes *BT3558* to *BT3564* as depicted in Figure 5A. In the seven other strains (jmh43, 60, 62, 63, 68, 71 and 72), the homology with VPI-5482 BT3563 was weaker (34% identity, Supplementary table S3), and the genetic environment of their *BT3563* homologs differed, including a homolog of *BT3559, BT3560, BT3564* as well as a gene coding for a protein carrying a *Bacteroides*-associated Carbohydrate binding Often N-terminal (BACON) domain Figure 5A. We also showed that *B. ovatus* and *B. uniformis* are the only Bacteroidales strains tested with a *BT3555-BT3564* region similar to *B. thetaiotaomicron* VPI-5482 including a homolog of *BT3560, BT3562* and *BT3563* (Supplementary figure S9). In the other strains tested, this region is either completely missing (*P. distasonis*) or present without a *BT3563* homolog (*B. fragilis, B. vulgatus, B. eggerthii)* (Supplementary figure S9). A more comprehensive search revealed that *BT3563* homologs were present in many diverse Bacteroidales, and at least in one example of *Flavobacterium columnare* and *Salinivirga cyanobacteriivorans,* non-Bacteroidales Bacteroidetes. However, the synteny of the *BT3560-3564* region was conserved only in strains from the *Bacteroides, Prevotella, Alloprevotella, Porphyromonas* and *Alistipes* genus (Supplementary figure S10). We tested the contribution of *BT3563* homologs to *B. thetaiotaomicron* bile-dependent biofilm formation in two randomly selected strains, jmh61 and jmh43. In these strains, the BT3563 proteins share respectively 87% and 34% of protein identity withVPI-5482 BT3563, and a different genetic organization around the *BT3563* homolog is observed (Supplementary Table S3 and Figure 5A). We showed that deletion of the *BT3563* homolog in both jmh61 and jmh43 led to a reduction of bile-dependent biofilm formation (Figure 5BC), indicating that the contribution of *VPI-5482* BT3563 extracellular DNase to bile-dependent biofilm formation is a widespread property of *B. thetaiotaomicron* species.

**Figure 5.**
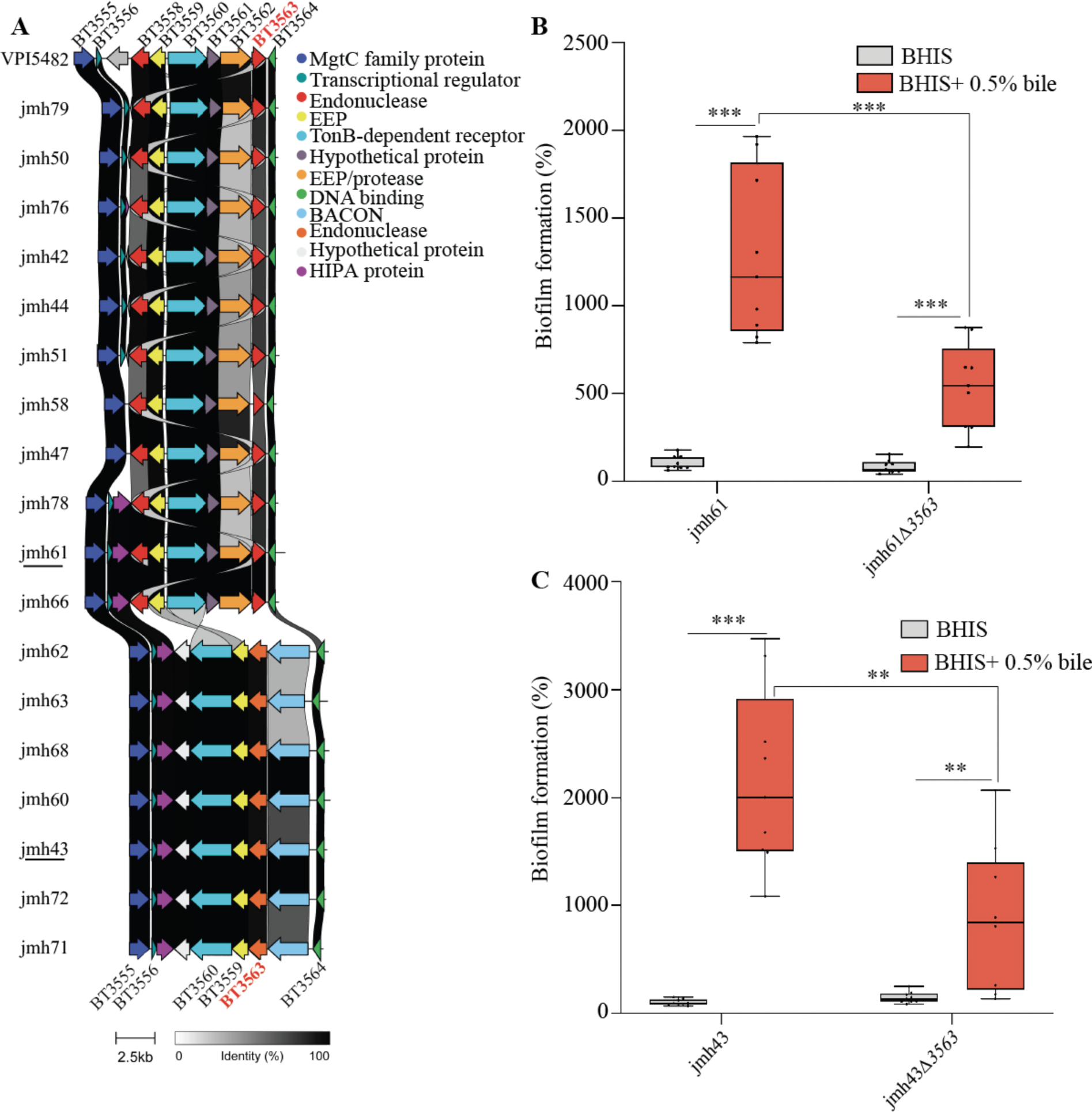
*BT3563* homologs are involved in bile-dependent biofilm formation of *B. thetaiotaomicron* clinical isolates. **A.** Comparison of the genetic organization around *BT3563* homologs in *B. thetaiotaomicron* isolates. EEP: endonuclease/exonuclease/phosphatase superfamily; BACON: *Bacteroides-*associated carbohydrate binding often N-terminal; HIPA: Serine/threonine-protein kinase toxin HipA. *BT3563* homologs sharing more than 80% sequence identity with VPI-5482 *BT3563* are shown in red (for jmh42, 44, 47, 50, 51, 58, 61, 66, 76, 78 and 79), *BT3563* homologs sharing 34% sequence identity with VPI-5482 *BT3563* are shown in orange (for jmh43, 60, 62, 63, 68, 71 and 72). **B.** and **C.** 96-well plate crystal violet biofilm assay after 48h growth in BHIS or BHIS+0.5% bile extract. Mean of jmh61 or jmh43 in BHIS is adjusted to 100 %. Min-max boxplot of 9 biological replicates for each strain, each replicate is the mean of two technical replicates. * p-value<0.05, ** p-value<0.005, *** p-value<0.0005, Mann-Whitney test.

## DISCUSSION

In this report, we showed that physiological concentrations of bile extract induce the formation of biofilm in almost all tested *B. thetaiotaomicron* strains, as well as in representative Bacteroidales species. This widespread induction by a compound exclusive to the intestinal environment suggests that biofilm formation might be an important aspect of Bacteroidales biology in the gut. Bile concentration in the gut can range between 0.2 to 2% (Kristoffersen et al., 2007), which is consistent with the concentrations required to induce maximum *B. thetaiotaomicron* biofilm formation (0.3%-1%). However, the concentration of intestinal bile is highly variable depending on the individual, time of the day, diet and the precise localization within the gut. Moreover, bile composition changes along the intestine and also depends on environmental parameters such as diet and gut microbiota composition. Thus, we can expect that the bile concentration and composition necessary to induce biofilm formation are only reached at certain times or in specific locations in the gut.

Although most studies have been performed with pathogenic bacteria, rather than with symbiotic members of the community, bile was previously shown to induce biofilm formation of at least one representative of all 4 major bacterial phylum of the gut microbiota: Firmicutes (*Clostridioides difficile, Listeria monocytogenes*), Bacteroidetes (*Bacteroides fragilis)*, Proteobacteria (*Vibrio cholerae)* and Actinobacteria (*Bifidobacterium)* (Ambalam et al., 2014; Begley et al., 2009; Dubois et al., 2019; Hung et al., 2006; Pumbwe et al., 2007), suggesting that bile is an important gut signal for biofilm formation. Consistently, we observed that bile extract induced biofilm formation in almost all tested *B. thetaiotaomicron* strains, as well as in representative strains of most of the common gut Bacteroidales species. Depending on the model considered in each of these studies, the concentration, incubation time and type of bile component necessary to trigger biofilm formation varied greatly. Bile-dependent biofilm formation also involved different ECM components, and different determinants. This suggests that although bile components might be a conserved signal for biofilm formation, the mechanism of biofilm induction differs. Bile could act as a signal triggering biofilm formation to stably colonize the gut. Alternatively, biofilm formation could be a conserved bacterial protection mechanism against the cytotoxic effects of bile, such as lipid membranes and DNA damage and protein denaturation (Urdaneta and Casadesús, 2017). Indeed, bacteria within biofilms are known to be more tolerant to various antimicrobials, including bile acids, than planktonic bacteria (Dubois et al., 2019; Hung et al., 2006; Pumbwe et al., 2007) and bile-dependent biofilm formation has been shown to increase tolerance to bile-mediated killing in different species including *B. fragilis* (Dubois et al., 2019; Hung et al., 2006; Pumbwe et al., 2007). Although, the production of a bile salt hydrolase (BSH) (Yao et al., 2018) and a tripartite multidrug resistance efflux pump, BT2793-2795 (Liu et al., 2019), confer a high resistance to bile to *B. thetaiotaomicron,* bile-induced biofilm formation in the gut probably constitutes an additional strategy by which gut bacteria protect themselves from bile toxicity (Flemming et al., 2016). Exposure of *B. thetaiotaomicron* to bile also induces various stress responses and increases the production of efflux pumps (Kristoffersen et al., 2007), suggesting the activation of stress tolerance mechanisms that could provide cross-protection against other damaging agents, such as antibiotics (Dubois et al., 2019; Pumbwe et al., 2007). Interestingly, in *Klebsiella pneumoniae,* an absence of poly-N-acetylglucosamine production significantly reduced bile-dependent biofilm formation *in vitro,* and also impaired its ability to colonize the mouse gut, showing that bile-dependent biofilm formation could impact colonization efficiency (Chen et al., 2014). Moreover, bile-mediated biofilm formation might be an important aspect of gut microbiota competition and cooperation. Bile is a complex mix of cholesterol, different bile acids and proteins. Some studies tested purified bile acids, rather than bile extract, and showed that all bacteria did not induce biofilm in response to the same bile acid. For instance, *C. difficile* reacted to deoxycholate (Dubois et al., 2019), *B. breve* to conjugated bile acids (taurocholate, glycocholate, taurodeoxycholate, glycodeoxycholate) (Kelly et al., 2020), and *Lactobacillus* to taurocholate (Ambalam et al., 2012). Conversely, taurine-conjugated bile acids disperse biofilms of *V. cholerae* or *P. aeruginosa* (Hay and Zhu, 2015; Sanchez et al., 2016). Interestingly, members of the gut microbiota can transform bile acids, with potential consequences on biofilm formation. In particular, several Clostridia can transform primary bile acids, such as cholate, into secondary bile acids, such as deoxycholate, by 7α-dehydroxylation and various bacteria possess a BSH enzyme that can remove the conjugated taurine or glycine residue of specific bile acids (Molinero et al., 2019). These bile modifications by members of the gut microbiota render bile acids less soluble and more cytotoxic, but they could also impact gut microbiota physiology (Cheng et al., 2019; Zhu et al., 2018) and the biofilm formation capacity of other bacteria, and therefore their colonization efficiency. Consistently, *Clostridium scindens* was shown to increase *C. difficile* biofilm formation *in vitro* by converting cholate to deoxycholate (Dubois et al., 2019).

Our results indicate that the simultaneous presence of bile, eDNA and the extracellular DNase BT3563 are required to observe dense, structured *B. thetaiotaomicron* biofilms. Presence of bile in the growth medium led to important physiological adaptations, as 17 % of *B. thetaiotaomicron* genes were differentially expressed, but the precise contribution of bile to biofilm formation is still unclear. Although the mechanism of bile-dependent biofilm formation has not been investigated for all bacteria, in some cases, bile was shown to increase the production of exopolysaccharide (Chen et al., 2014; Crawford et al., 2008; Nickerson et al., 2017), cyclic-di-GMP (Koestler and Waters, 2014) and various adhesins such as autotransporters (Köseoglu et al., 2019) and curli (González et al., 2019). However, very little is known about *B. thetaiotaomicron* biofilm determinants. Capsular polysaccharide 8 and type V pili production were previously shown to increase *B. thetaiotaomicron* adhesion capacity, but these genes were not induced in the presence of bile. However, *B. thetaiotaomicron* mucin O-glycan degrading PULs were up-regulated in biofilms grown in chemostats compared to planktonic cultures, and in the mucus compared to the lumen (Li et al., 2015; TerAvest et al., 2014) and they were also induced in presence of bile, suggesting bile exposure might prime bacteria for adhesion and colonization to the mucus layer *in vivo.* Bile also increased the release of eDNA, which could modulate biofilm formation. eDNA is a well-known component of the biofilm ECM and numerous studies have shown that addition of DNase to the growth media prevented biofilm formation or dispersed pre-formed biofilm in both Gram-positive and Gram-negative bacteria (Okshevsky and Meyer, 2015). Consistently, addition of DNase I led to a small reduction in the bile-dependent biofilm formation of *B. thetaiotaomicron,* suggesting that eDNA is an important ECM component (Figure 3B). However, unexpectedly, we show that the extracellular DNase BT3563 is also necessary for bile-dependent biofilm formation, suggesting that partial degradation of eDNA might be necessary for biofilm formation in presence of bile. Neither *BT3563* transcription nor BT3563 DNase activity was increased by bile, so the link between bile exposure and BT3563 function remains unclear. BT3563 carries a lipoprotein signal commonly found in surface-exposed outer membrane proteins in *Bacteroides.* Consistently, BT3563 has been detected in the *B. thetaiotaomicron* outer-membrane and was enriched in outer membrane vesicles (OMV) (Valguarnera et al., 2018), suggesting BT3563 might be secreted to the supernatant through OMV production. We hypothesize that cell- or OMV-associated BT3563 could degrade eDNA during biofilm formation, maintaining a specific eDNA concentration or organization within the ECM to form structured bile-dependent biofilms. Deletion of *BT3563* led to the formation of scattered biofilms in presence of bile, composed of clumps of cells rather than a dense layer at the bottom of the well. To allow the formation of denser structures, BT3563 might be required to remove eDNA from cell vicinity and prevent electrostatic repulsion between the negatively charged eDNA and bacterial surface or some component of the ECM such as exopolysaccharides. For instance, the production of positively-charged proteins was necessary to prevent electrostatic repulsion between eDNA and *Staphylococcus aureus* cells in biofilms and perturbation of these proteins led to an increasingly porous biofilm, reminiscent of our *ΔBT3563* biofilms (Dengler et al., 2015; Kavanaugh et al., 2019). It is also to be noted that a *ΔBT3560-3563* mutant formed less bile-dependent biofilm than a *ΔBT3563* simple mutant, suggesting the TonB-dependent receptor BT3560 and/or the nuclease BT3562 could also contribute to biofilm formation. Moreover, *ΔBT3560-3563* still formed biofilm in the presence of bile, demonstrating that additional factors could be involved.

Our study therefore demonstrates that gut-relevant environmental cues such as bile can strongly stimulate *B. thetaiotaomicron* biofilm formation. This induction relied on the production of an extracellular DNase, BT3563, that is present in all *B. thetaiotaomicron* isolates, and in many diverse Bacteroidetes. Although the contribution of these BT3563 homologs to bile-dependent biofilm formation will need to be confirmed beyond the three isolates tested in this study, this study suggests that, in addition to the well-established structural role of eDNA in the biofilm matrix, partial eDNA degradation might also represent an important aspect of biofilm formation in *B. thetaiotaomicron* species, and possibly in other Bacteroidales.

## MATERIALS AND METHODS

### Bacterial strains and growth conditions

Bacterial strains used in this study are listed in Supplementary Table S4. *B. thetaiotaomicron* and other Bacteroidales strains were grown in BHIS broth (Bacic and Smith, 2008) supplemented with erythromycin 15 µg/ml (erm), tetracycline 2.5 µg/ml (tet), gentamycin 200 µg/ml (genta), 5’-fluoro-2’-deoxyruidin 200 µg/ml (FdUR), anhydrotetracycline (0.1 µg/mL), D-glucose, D-mannose, D-rhamnose, D-cellobiose, D-maltose (0,5% w/v), hemin from bovine (25mg/L and 50mg/L), porcine mucin extract (0,1% and 0,5% w/v), bile extract from bovine and ovine (Sigma, B8381) at 0.5% unless indicated otherwise, DNase I (Thermo scientific, VF304452) 98U/mL, or RNase1 (Thermo Scientific, EN0601) 0.06U/mL when required. Cultures were incubated at 37°C in anaerobic conditions using jars with anaerobic atmosphere generators (GENbag anaero, Biomerieux, ref. 45534) or in a C400M Ruskinn anaerobic-microaerophilic station. *Escherichia coli* S17λpir was grown in Miller’s Lysogeny Broth (LB) (Corning) supplemented with ampicillin (100 µg/ml) when required and incubated at 37°C with 180 rpm shaking. Cultures on solid media were done in BHIS with 1.5% agar and antibiotics were added when needed. Bacteria were always streaked from glycerol stock on BHIS-agar before being grown in liquid cultures. All media and chemicals were purchased from Sigma-Aldrich unless indicated otherwise.

All experiments and genetic constructions of *B. thetaiotaomicron* were made in VPI-5482*Δtdk* background (Koropatkin et al., 2008) unless indicated otherwise, which was developed for 2-step selection procedure of unmarked gene deletion by allelic exchange, as previously described. Therefore, the VPI-5482*Δtdk* is referred to as wild type or VPI-5482 in this study.

### Genome analysis

Genomic DNA of clinical strains was prepared from overnight cultures using the DNeasy blood and tissue kit (Qiagen). Illumina whole genome sequencing was performed by the Plateforme de microbiologie mutualisée (P2M) of Institut Pasteur and the genomes were assembled SPAdes v3.13.0 (Bankevich et al., 2012), and when necessary were reassembled using Unicycler (Wick et al., 2017). The obtained genomes were annotated using RASTtk (Brettin et al., 2015; Wattam et al., 2017) on the patricbrc.org database (Davis et al., 2020) and we searched *BT3563* nucleic acid and amino acid sequence in these genomes using the BLASTp tool of patricbrc.org (Boratyn et al., 2013; O’Leary et al., 2016). All of these genomes were made available publicly on the patricbrc.org database (see supplementary table S1 for accession numbers). Moreover, this Whole Genome Shotgun project has been deposited at DDBJ/ENA/GenBank under the Bioproject accession PRJNA725886. The accession numbers for each strain are shown in supplementary table S1. Sequences of the region surrounding *BT3563* homologs were extracted and the gene cluster comparison was made using the clinker and clustermap.js pipeline (Gilchrist and Chooi, 2021). Synteny analysis was performed using either the web-based tool Synttax (Oberto, 2013) searching for homologs of *BT3563* using a custom selection of diverse Bacteroidetes, or the genome bowser tool of the MicroScope plateforme (Vallenet et al., 2020) looking for *B. thetaiotaomicron BT3555-3564* region homology in the seven Bacteroidales strains tested.

### 96-well crystal violet biofilm formation assay

Overnight culture was diluted to OD_600_ = 0.05 in 100 µL BHIS and inoculated in technical duplicates in polystyrene Greiner round-bottom 96-well plates. The wells at the border of the plates were filled with 200 µL of water to prevent evaporation. Incubation was done at 37°C in anaerobic conditions for 48h. The supernatant was removed by careful pipetting and the biofilms were fixed using 100 µL of Bouin’s solution (picric acid 0.9%, formaldehyde 9% and acetic acid 5%, HT10132, Sigma-Aldrich) for 10min. Then the wells were washed once with water by immersion and flicking, and the biofilm was stained with 125 µL 1% crystal violet (V5265, Sigma-Aldrich) for 10 minutes. Crystal violet solution was removed by flicking and biofilms were washed twice with water. Stained biofilms were resuspended in 1:4 acetone: ethanol mix and absorbance at 575 nm was measured using TECAN infinite M200 PRO plate reader.

### Targeted mutagenesis

A list of all the primers used in this study can be found in Supplementary Table S5. Deletion mutants in *B. thetaiotaomicron* VPI 5482 were constructed using the previously described vector for allelic exchange pExchange*-tdk* (Koropatkin et al., 2008). Deletion mutants in *B. thetaiotaomicron* clinical isolates were constructed using the recently described suicide vector pLGB13 for allelic exchange in *Bacteroides* natural isolates (Addgene 126618) (García-Bayona and Comstock, 2019). Briefly, 1kb region upstream and downstream of the target sequence and the vector (pExchange-*tdk* or pLGB13) were amplified by PCR using Phusion Flash High-Fidelity PCR Master Mix (Thermofischer Scientific, F548). All three fragments were ligated using Gibson assembly: the inserts and the plasmids were mixed with Gibson master mix 2x (100 µL 5X ISO Buffer, 0.2 µL 10,000 U/mL T5 exonuclease (NEB #M0363S), 6.25 µL 2,000 U/mL Phusion HF polymerase (NEB #M0530S), 50 µL 40,000 U/mL Taq DNA ligase (NEB #M0208S), 87 µL dH2O for 24 reactions) and incubated at 50°C for 35 min. The resulting mix was transformed in *E. coli* S17λpir that was used to deliver the vector to *B. thetaiotaomicron* by conjugation. Conjugation was carried out by mixing exponentially grown cultures (OD_600_=0.6) of the donor and the recipient strain in a 2:1 ratio. The mixture was spotted on BHIS-agar plates and incubated at 37°C in aerobic conditions overnight. The mix was then streaked on BHIS agar supplemented with antibiotic – for selection of *B. thetaiotaomicron* transconjugants that had undergone the first recombination event – and gentamicin to ensure exclusion of any *E. coli* growth. Eight of the resulting colonies were grown overnight in BHIS with no antibiotic to allow a second recombination event, and the culture was plated on BHIS-agar plates supplemented with either FdUR (to counter select pEchange-*tdk*) or anhydrotetracycline (to counter select pLGB13) to select for loss of plasmid. The resulting deletion mutants were confirmed by PCR and sequencing.

We used the pNBU2-bla-erm vector (Wang et al., 2000) or pNBU2-bla-tet (Béchon et al., 2020b) vector for complementation, which inserts in the 5’ untranslated region of the tRNA-Ser, in which we previously cloned the constitutive promoter of *BT1311* encoding the sigma factor RpoD (Mihajlovic et al., 2019). Target genes were amplified by PCR using Phusion Flash High-Fidelity PCR Master Mix from start codon to stop codon and they were cloned after *BT1311* promoter by Gibson assembly. The Gibson mix was transformed in *E. coli* S17λpir and the resulting *E. coli* was used to transfer the plasmid to *B. thetaiotaomicron* by conjugation (see above).

### Random transposon mutagenesis

pSAMbt, the previously published tool for random mariner-based transposon mutagenesis in *B. thetaiotaomicron* (Goodman et al., 2009) was conjugated in *B. thetaiotaomicron* as described above. After streaking on BHIS-erm-genta agar plates, isolated colonies were resuspended in 100 µL BHIS in 96-well plates, grown overnight and tested for biofilm formation as described above. The selected clones were then streaked on a fresh BHIS-erm-genta agar plate and 3 isolated colonies were tested for biofilm formation to ensure no mix of transposon mutants had occurred during preparation of the library. The genomic DNA of the validated clones was extracted using DNeasy blood and tissue kit (Qiagen) and sent for whole genome sequencing at the Mutualized platform for Microbiology of Institut Pasteur.

### Growth curve

2.5 µL overnight cultures were added to 200µL BHIS that had previously been incubated in anaerobic condition overnight to remove dissolved oxygen, in Greiner flat-bottom 96-well plates. A plastic adhesive film (adhesive sealing sheet, Thermo Scientific, AB0558) was added on top of the plate inside the anaerobic station, and the plates were then incubated in a TECAN Infinite M200 Pro spectrophotometer for 24 hours at 37°C. OD600 was measured every 30 minutes, after a 900-second orbital shaking of 2 mm amplitude.

### Viable cells count

Overnight cultures were diluted to OD_600_ = 0.5 in a final volume of 1mL of NaCl 0.85%. Aliquots were centrifugated at 6000 g (7000rpm) for 7 min and the supernatant was discarded. After washing twice with NaCl 0.85%, 2µL of propidium iodide was added and the samples were incubated in anaerobic condition and protected from light at 37°C, for 30 minutes. The number of viable bacteria was then quantified using MACSQuant VYB flow cytometer and MACSQuantify Software V2.11.

### Extracellular matrix extraction and quantification

ECMs were extracted based on previously described methods (Chiba et al., 2015). 2mL of 48h biofilms cells grown in presence of bile were harvested and centrifugated at 5000 × g for 10 min. The pellet was washed twice with NaCl 0.85% and weighed, then resuspended in an extraction buffer (Tris-HCl pH 8.0; 1.5 M NaCl) at a 1:10 mass-volume ratio and incubated at 25 °C for 30 min with agitation. Then, cells were removed by centrifugation at 15,000 × g and 25 °C for 10 min and the supernatant containing the extracted ECMs were stored at −20°C until use. The amount of RNA, DNA and proteins in the ECM were measured using a Qubit 3.0 Fluorometer (Thermo Fisher Scientific) according to the manufacturer’s instructions.

### Nuclease activity of the supernatant

Overnight cultures were centrifugated 6.5 min at 6000 x g. When applicable, we measured eDNA concentration using the QuBit HS double stranded DNA kit (Invitrogen). 50 µl of supernatant were mixed with *B. thetaiotaomicron* VPI-5482 genomic DNA. 10 µL were used to run on a 0.8% agarose gel and colored with ethidium bromide. The remaining 40 µL were incubated at 37°C overnight for approximatively 18h and then 10 µL were used to run a 0.8% agarose gel and colored with ethidium bromide.

### Confocal Laser Scanning Microscopy (CLSM)

For confocal laser scanning microscopy, biofilms were grown in 96-well plates (µclear, Greiner Bio-One). 100 µL of BHIS, supplemented with 0.5% bile extract, DNase I (Thermo scientific, VF304452) 98 U/mL or RNase1 (Thermo Scientific, EN0601) 0.06 U/mL when required, was added to each well and the plates were incubated at 37 °C, in static condition 48h under anaerobic conditions. The unwashed biofilms were then directly stained in red with 20 μM of SYTO61 (Life Technologies; cell permeant nucleic acid dye to contrast all the bacteria) and in green with 0.4 µM of TOTO-1 (Thermoscientific; cell impermeant DNA dye to contrast eDNA). After 15 min of incubation, Z-stacks of horizontal plane images were acquired in 1 μm steps using CLSM (Leica TCS SP8, INRAE MIMA2 microscopy platform) with a water 63× immersion lens (NA = 1.2). Two stacks of images were acquired randomly on three independent samples at 800 Hz. Fluorophores were excited and emissions were captured as prescribed by the manufacturer. Simulated 3D fluorescence projections were generated using IMARIS 9.3 software (Bitplane, Zürich, Switzerland).

### RNAseq analysis

Overnight cultures were mixed with RNAprotect (Qiagen) to prevent RNA degradation, and the pellet was kept at −80°C. Total RNA was extracted using MP Biomedicals FastRNA Pro^TM^ BLUE KIT by the provider’s manual and treated with Ambion Turbo DNA-free™ kit to remove possible DNA contamination. Total RNA from 4 independent replicates were checked on RNA6000 RNA chips on Bioanalyzer (Agilent) for its quality and integrity. Ribosomal RNA depletion was performed using the Bacteria RiboZero kit (Illumina). From rRNA-depleted RNA, directional libraries were prepared using the TruSeq Stranded mRNA Sample preparation kit following the manufacturer’s instructions (Illumina). Libraries were checked for quality on Bioanalyzer DNA chips (Agilent). Quantification was performed with the fluorescent-based quantitation Qubit dsDNA HS Assay Kit (Thermo Fisher Scientific). Sequencing was performed as a Single Read run for 65 bp sequences on HiSeq 2500 Illumina sequencer (65 cycles). The multiplexing level was 16 samples per lane. Bioinformatics analysis were performed using the RNA-seq pipeline from Sequana.

## ACKNOWLEDGEMENTS

We thank Rebecca Stevick, Christophe Beloin, Susan Joyce and David Clarke for critical reading of the manuscript. We are grateful to Andy Goodman, Justin Sonnenburg and Laurie Comstock for providing the genetic tools used in this study. This work was supported by an Institut Pasteur grant and by the French government’s Investissement d’Avenir Program, Laboratoire d’Excellence “Integrative Biology of Emerging Infectious Diseases” (grant n°ANR-10-LABX-62-IBEID) and the *Fondation pour la Recherche Médicale* (grant no. DEQ20180339185). N. Béchon was supported by a MENESR (Ministère Français de l’Education Nationale, de l’Enseignement Supérieur et de la Recherche) fellowship. J. Mihajlovic was supported by the Pasteur Paris University International Doctoral Program and the *Fondation pour la Recherche Médicale* (grant no. FDT20160435523). The RNAseq experiments was performed by the Biomics Platform, C2RT, Institut Pasteur, Paris, France, supported by France Génomique (ANR-10-INBS-09-09) and IBISA.

## AUTHOR CONTRIBUTIONS

N.B., J.M. and J.-M.G. designed the experiments. N.B. performed the bioinformatic and sequence analyses. N.B., J.M., A.A.L. and S.V.-F. performed the experiments. N.B., J.M., A.A.L. and J.-M.G. analyzed data. B.D. and I. M.-V. contributed to the initial experiments and provided advice. O.S. performed RNAseq analysis, J.D. and R. B. performed the image analyses. J.-M.G, N.B., and J.M. wrote the paper with significant contribution from all co-authors.

## SUPPLEMENTARY MATERIALS

### SUPPLEMENTARY FIGURES

**Supplementary Figure S1.**
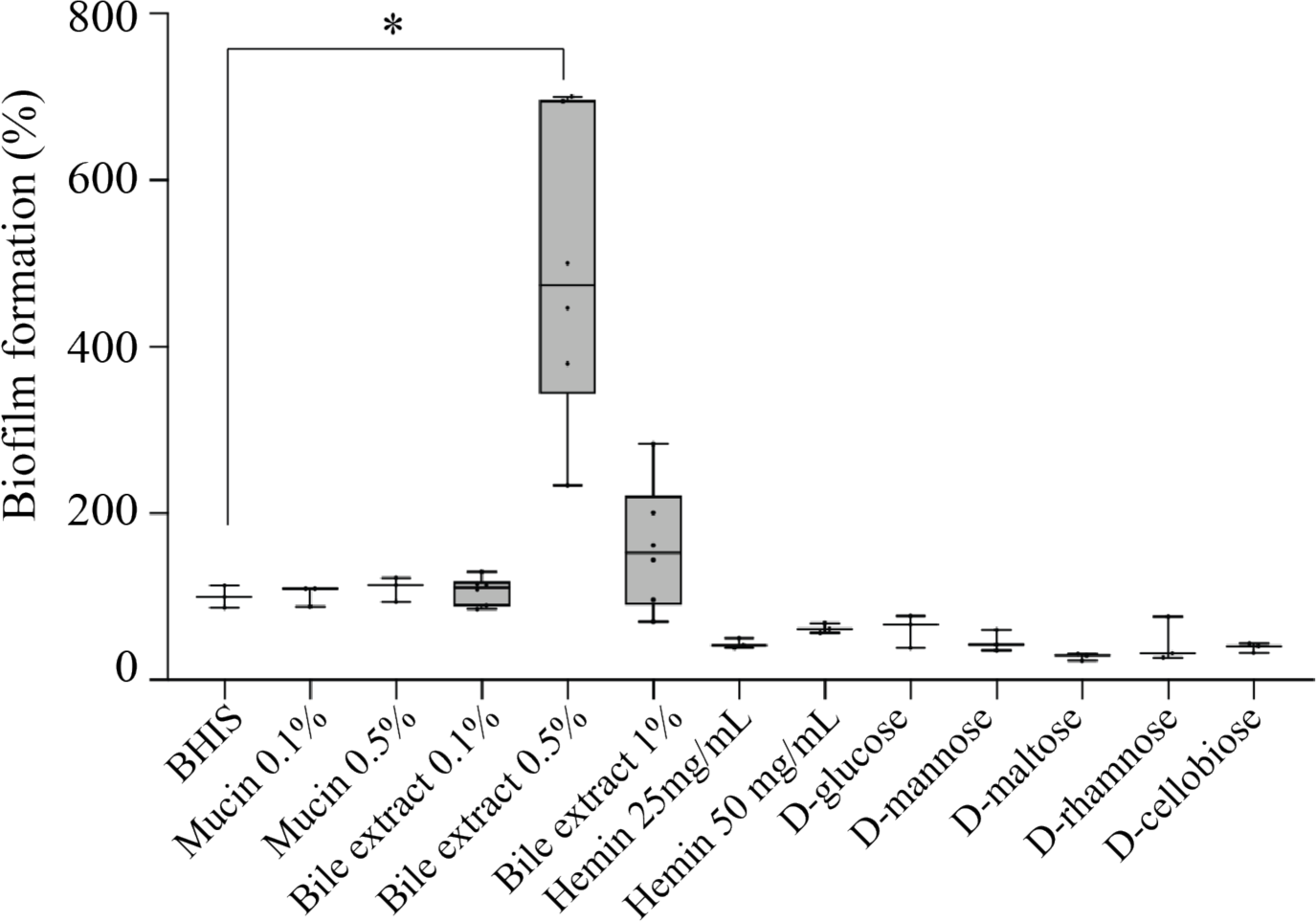
Bile is the only tested environment cue that induces biofilm formation of *B. thetaiotaomicron* VPI-5482. *B. thetaiotaomicron* VPI-5482 after 24h growth in BHIS supplemented with 0.5% D-glucose, D-mannose, D-rhamnose, D-cellobiose and D-maltose, 25 mg/L and 50 mg/L hemin, 0.1% and 0.5% mucin, and 0.1%, 0.5%, 1% bovine-ovine bile extract. Min-max boxplot of 3-6 biological replicates for each strain. * p-value<0.05, ** p-value<0.005, Mann-Whitney test.

**Supplementary Figure S2.**
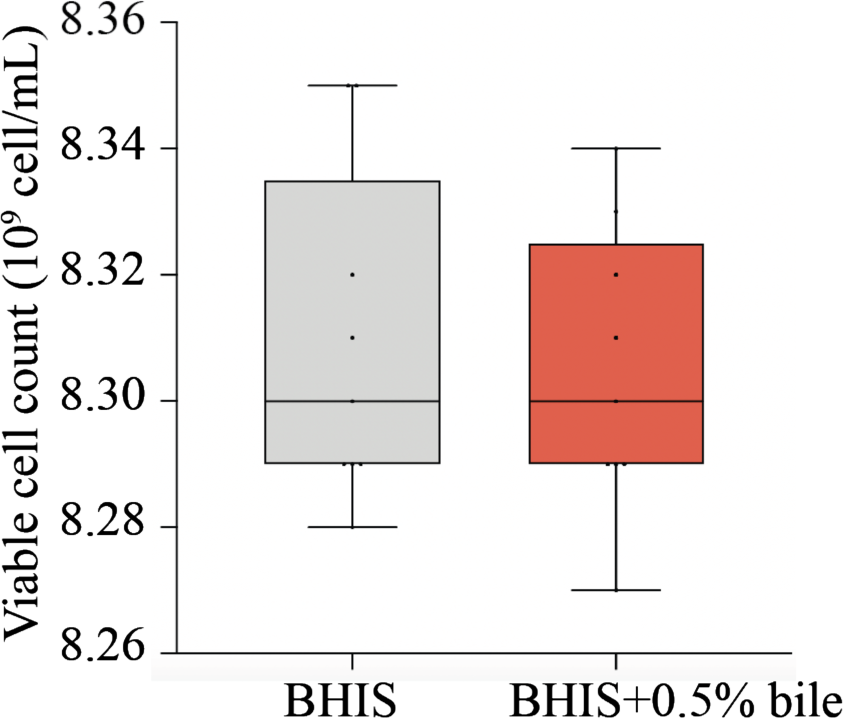
0.5% bile extract concentration is non-toxic for *Bacteroides thetaiotaomicron*. **A.** Flow cytometry of *B. thetaiotaomicron* VPI-5482 cultures grown in presence or absence of bile and dyed with propidium iodide to label live cells. The viable cell count is shown as a min-max boxplot of 6 biological replicates.

**Supplementary Figure S3.**
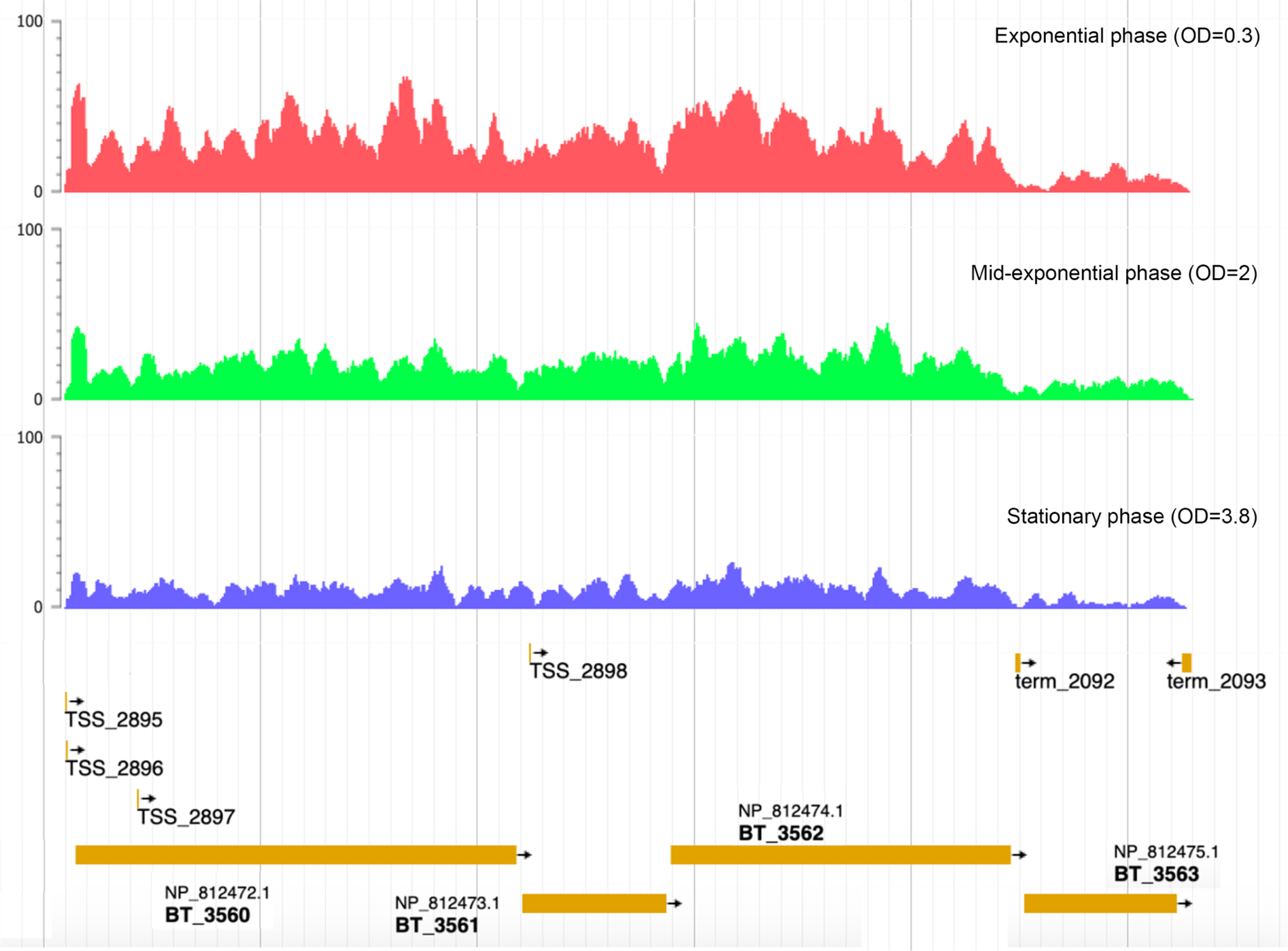
*BT3560-BT3563* operon. RNAseq reads from *B. thetaiotaomicron* VPI-5482 grown in TYG medium to exponential phase (OD=0.3) in red, mid-exponential phage (OD=2) in green and stationary phase (OD=3.8) in blue are mapped to the *BT3560-BT3563* genetic locus. Predicted transcriptional start sites (TSS) and terminators (term) are depicted, showing the predicted operon structure of *BT3560-BT3563*. Data and bioinformatics prediction from Theta-Base database (Ryan et al., 2020) (https://bacteroides.helmholtz-hzi.de/) plotted on JBrowser (Buels et al., 2016).

**Supplementary Figure S4.**
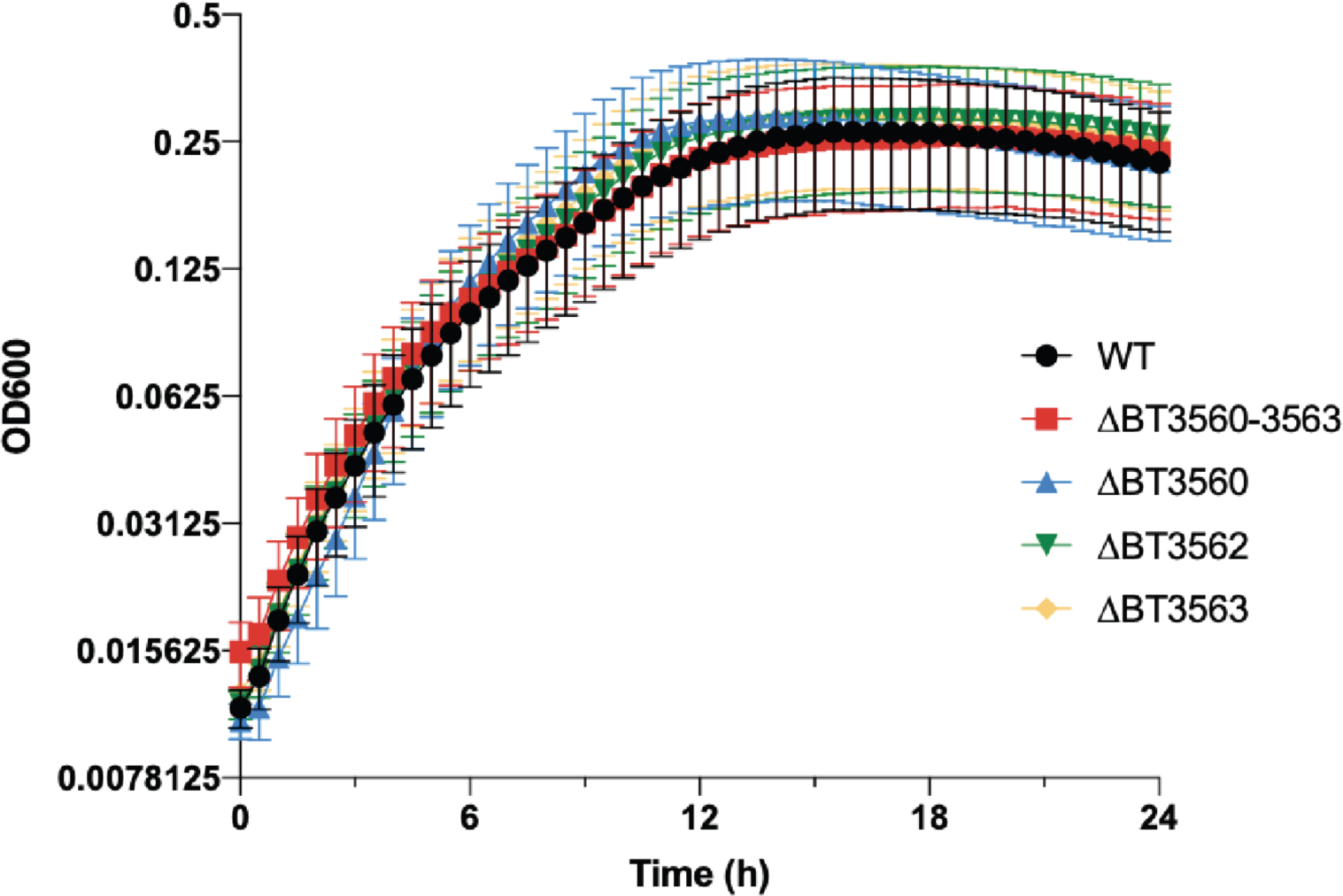
Growth curve. 24h growth curve of indicated strains in 96-well plate in BHIS+0.5% bile extract. Mean of 6 biological replicates, error bars represent standard error to the mean (SEM).

**Supplementary Figure S5.**
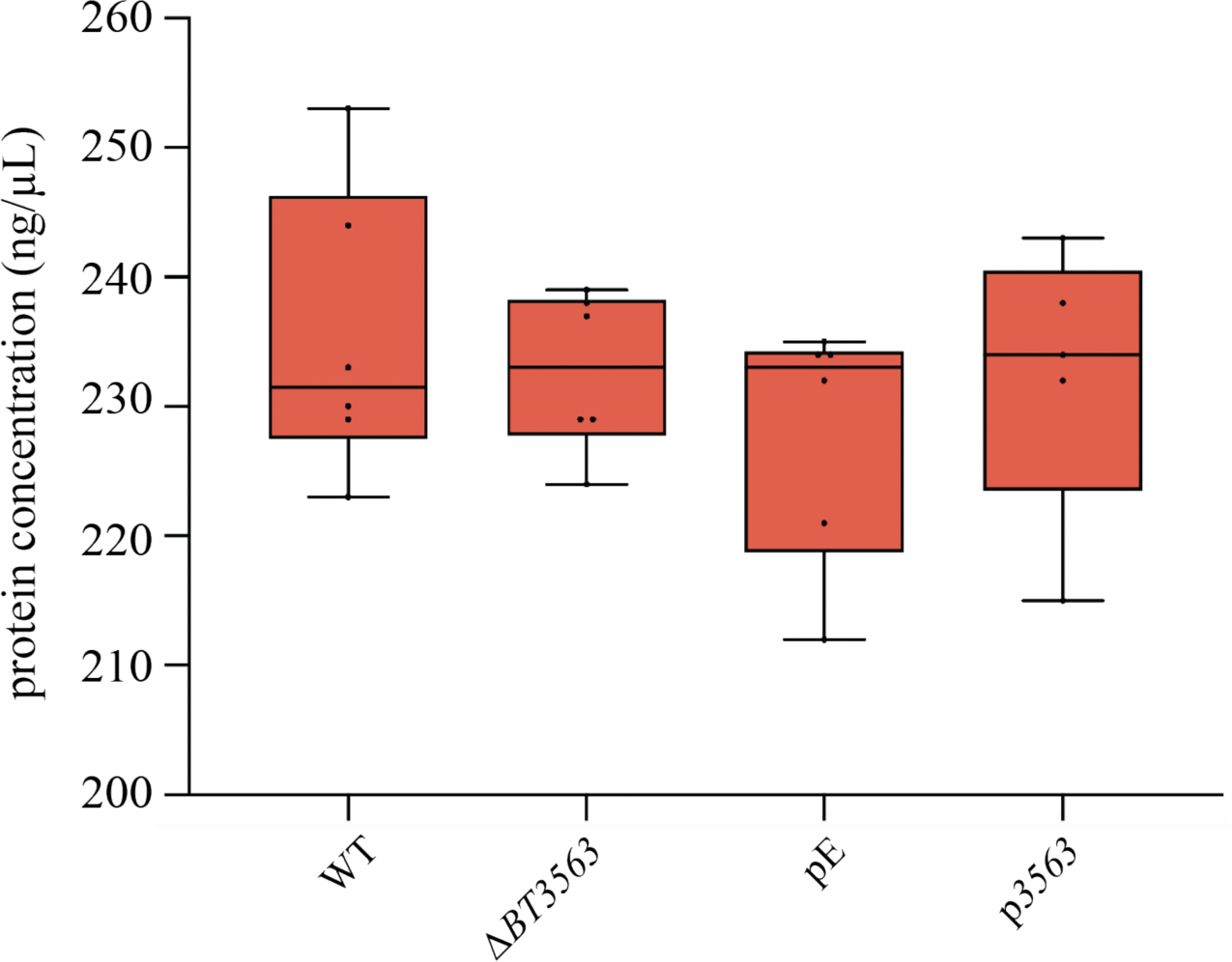
BT3563 does not impact protein concentration in the extracellular matrix. Extracellular protein concentration (ng/µL) in the purified extracellular matrix of bile-dependent biofilms grown in BHIS+0.5% bile extract for 48h. Min-max boxplot of 6 biological replicates for each strain. * p-value<0.05, ** p-value<0.005, Mann-Whitney test.

**Supplementary Figure S6.**
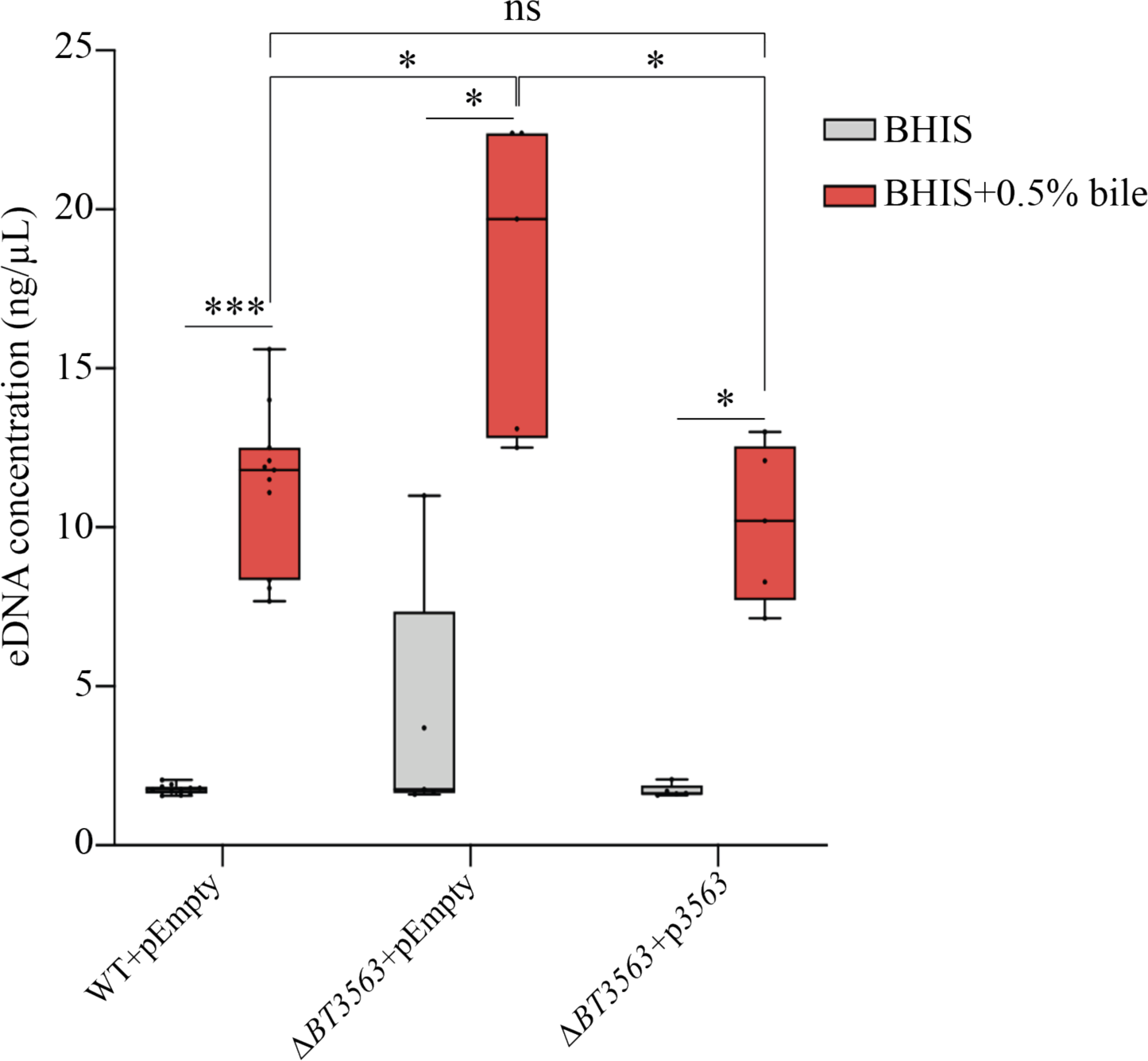
Bile increases eDNA concentration in the supernatant. Extracellular DNA concentration (ng/µL) in the supernatant of overnight cultures of indicated strains grown in BHIS or BHIS+0.5% bile extract. Min-max boxplot of 6-11 biological replicates for each strain. * p-value<0.05, *** p-value<0.0005, Mann-Whitney test.

**Supplementary Figure S7.**
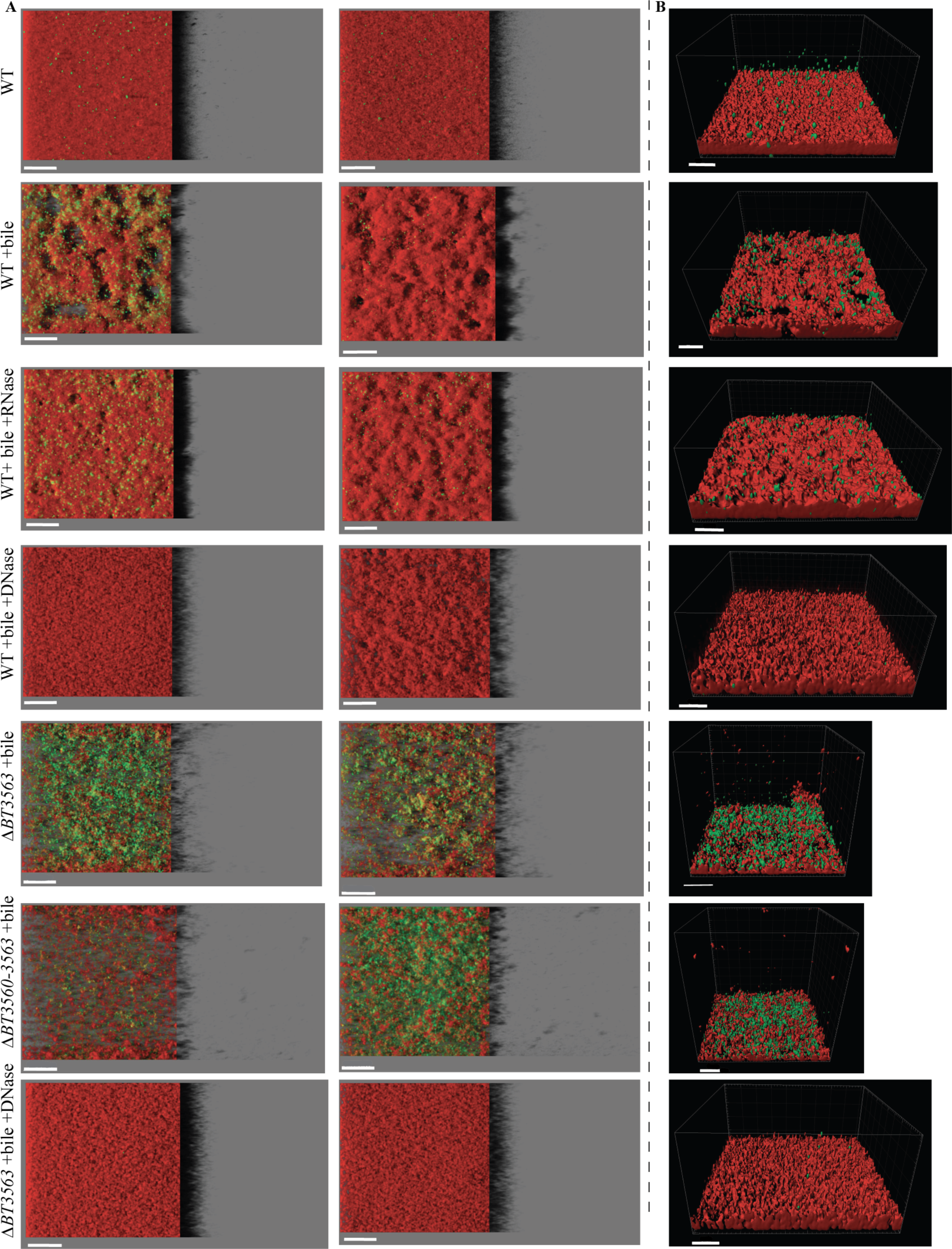
Visualization of *B. thetaiotaomicron* biofilms. **A.** and **B.** *B. thetaiotaomicron* biofilms grown in plates in absence or presence of 0.5% bile extract (“+bile”), RNase I (“+RNase”), or DNase I (“+DNase”). Merged images showing cells labelled with SYTO61 dye in red and extracellular DNA and dead cells labelled with TOTO-1 in green. **A.** IMARIS easy 3D projections of CLSM images of two additional wells of *B. thetaiotaomicron* biofilms grown in plates. For each image, the shadow projection of the biofilm is shown on the right. Scale bar represent 40 µm. **B.** A 3D reconstruction of *B. thetaiotaomicron* biofilm. Scale bar represent 30 µm.

**Supplementary Figure S8.**
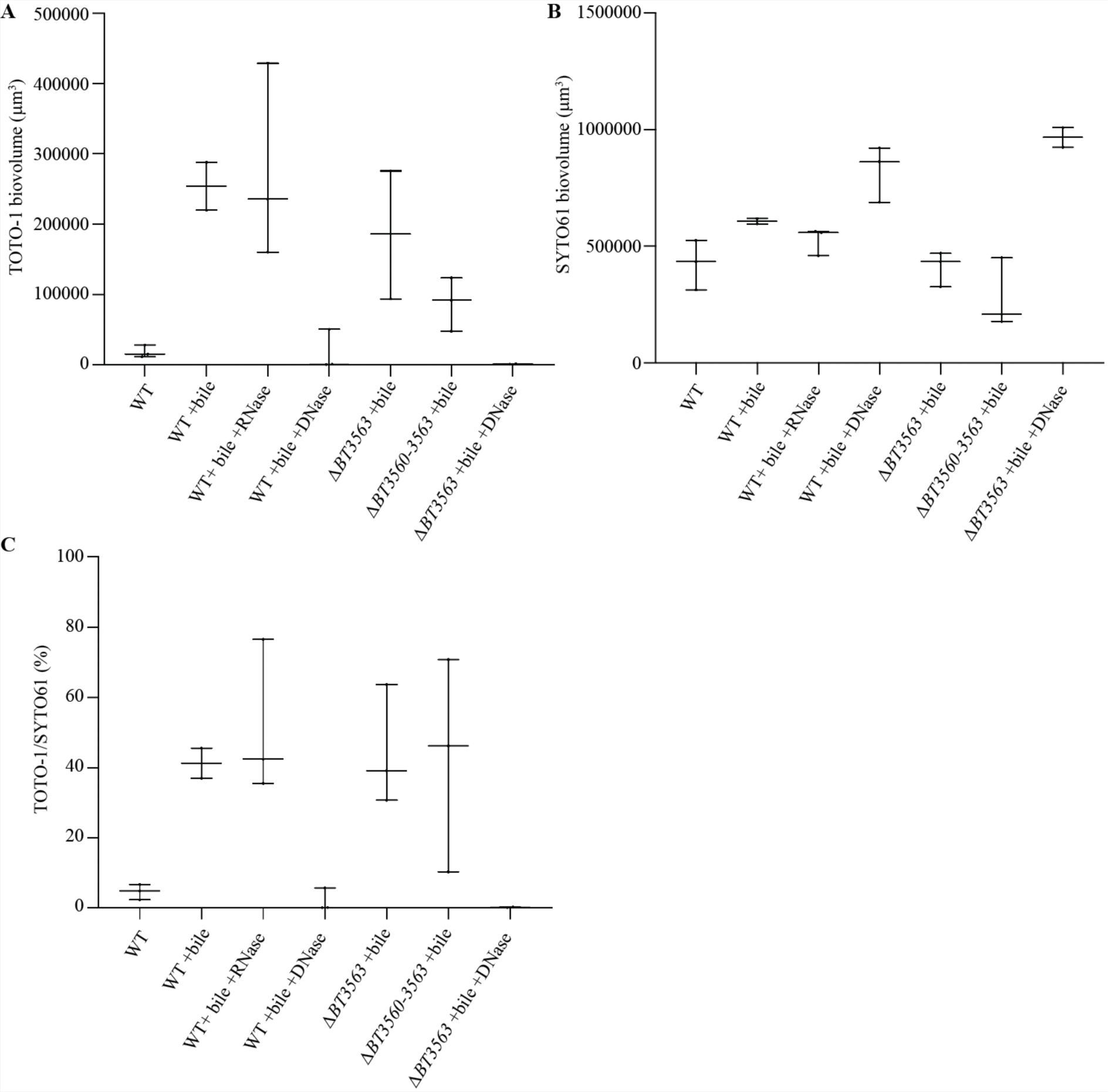
Quantification of confocal microscopy images. Min-Max boxplot of **A.** biovolumes of TOTO-1, of **B.** SYTO61 fluorescence (µm^3^) and **C.** TOTO-1/SYTO61 fluorescence ratio (%). 2-3 biological replicates, each representing the mean of 2-3 technical replicates.

**Supplementary Figure S9.**
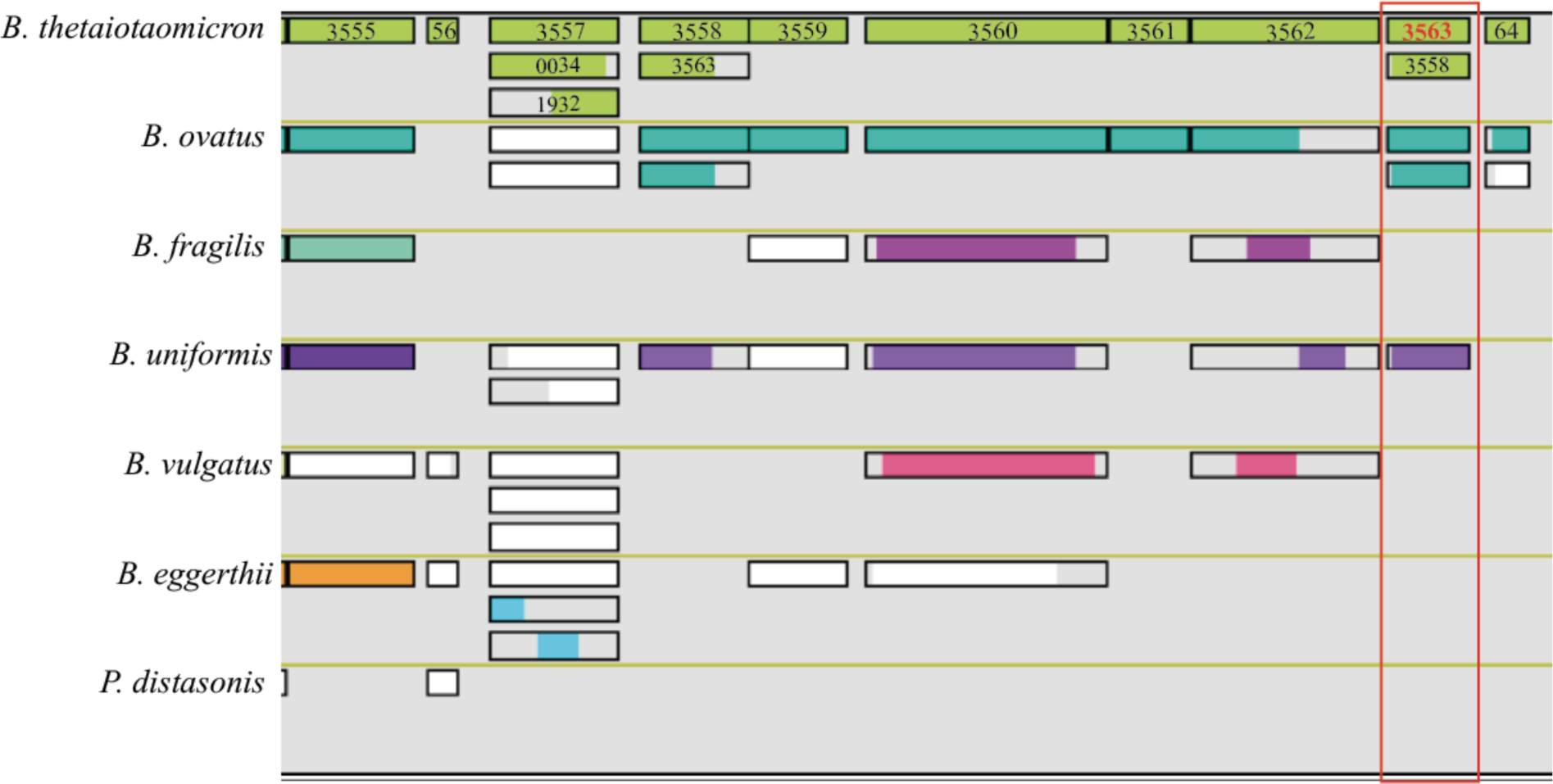
*BT3555-3564* synteny among the tested Bacteroidales. Synteny analysis was performed using the Genome browser tool of the genome analysis platform MicroScope, (https://mage.genoscope.cns.fr/microscope/mage/viewer.php?) (Vallenet et al., 2020). For each genome, the first line represents the genes on the chromosome, and the additional lines represent homologs of the considered gene within the same genome, when applicable. The colored bar represents the length of the homology. Locus tags are indicated for *B. thetaiotaomicron* VPI-5482 strain, and the homologs of *BT3563* are indicated in a red square.

**Supplementary Figure S10.**
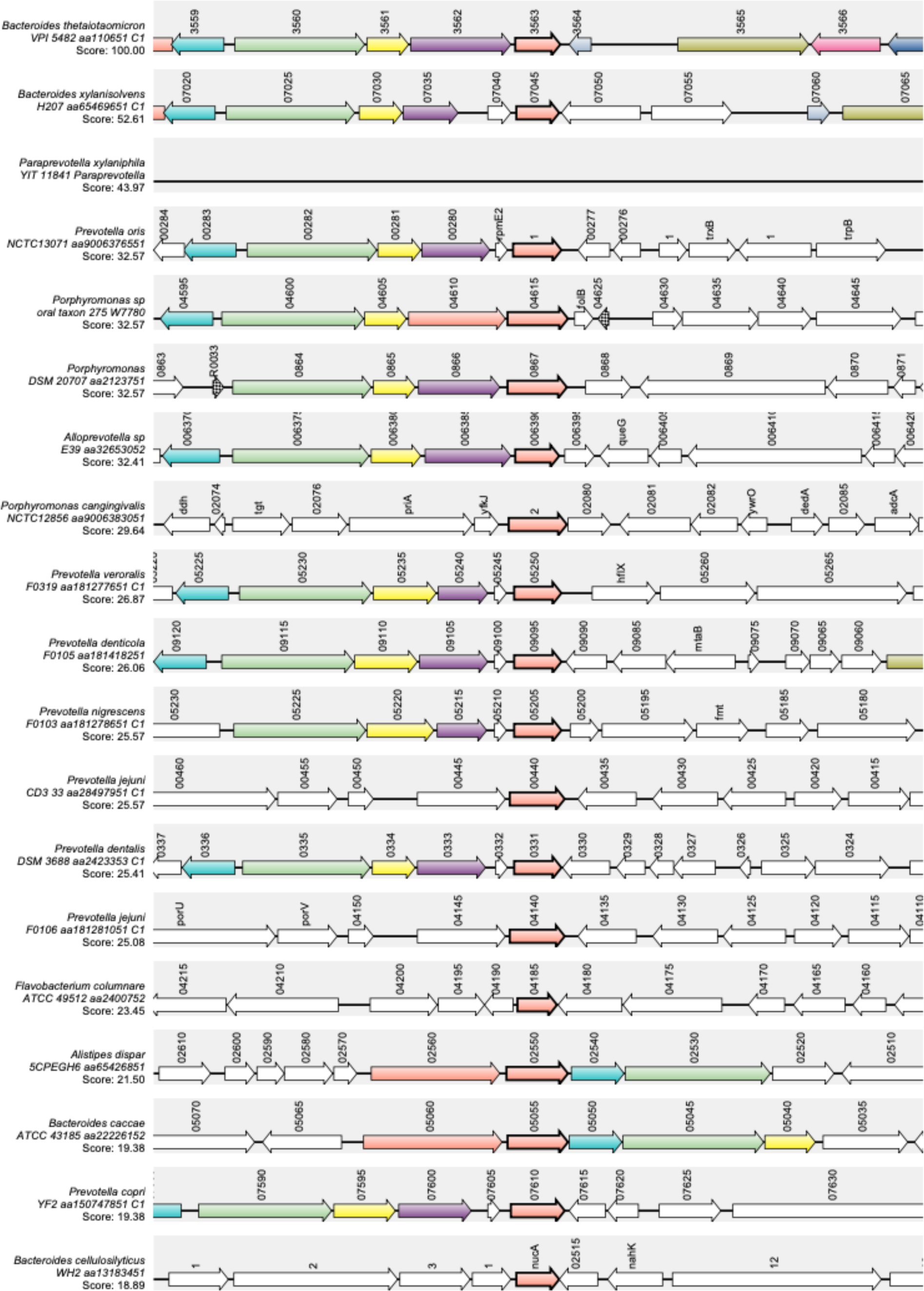
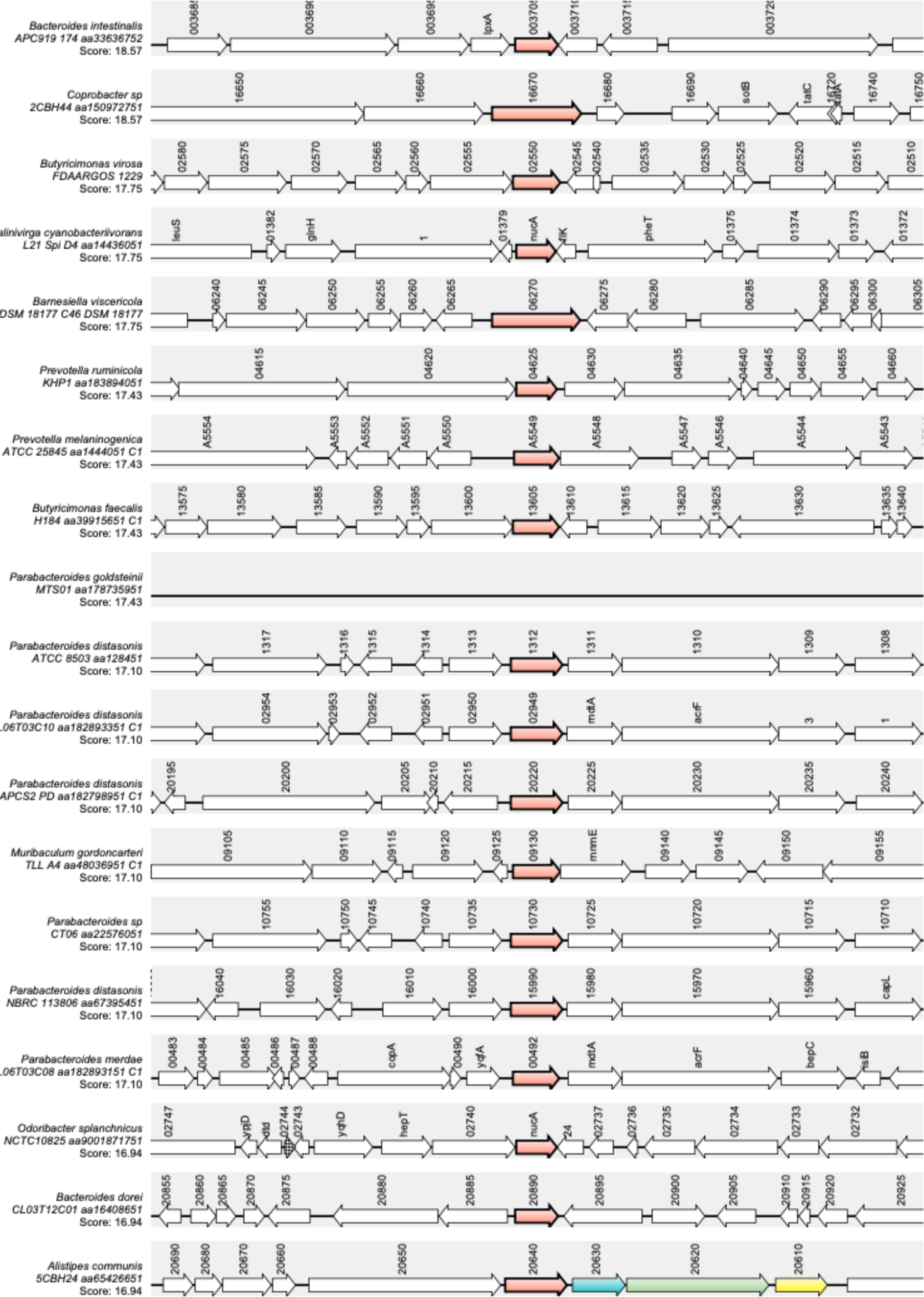
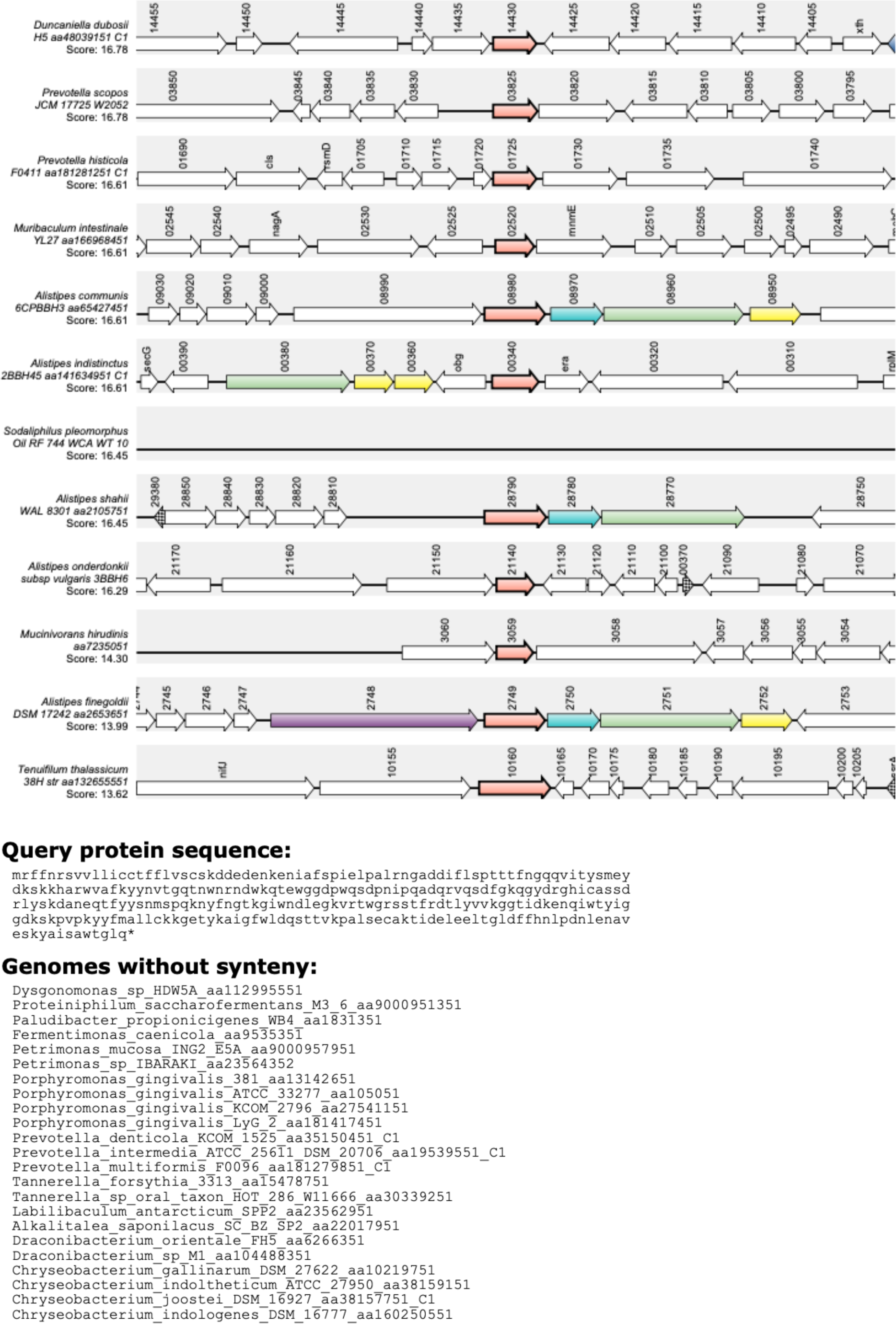
*BT3563* synteny among diverse Bacteroidetes. Synteny analysis was performed using the web tool SyntTax (Oberto, 2013). Homologs of the gene of interest, *BT3563,* are marked in bold. Homologous genes are represented in the same color. Locus tags are annotated on top of each gene. A list of all tested strains for which no *BT3563* homologs was found is present at the bottom.

### SUPPLEMENTARY TABLES

**Supplementary Table S1.** *Bacteroides thetaiotaomicron* strains used in this study

| Strain | Strain name 2 | Origin | From | PATRIC ID | Genbank ID |
| --- | --- | --- | --- | --- | --- |
| jmh42 | N03576 | Perforated ulcer | CHU Limoges | 818.1412 | JAGURM000000000 |
| jmh43 | N03525 | Blood | CHU Limoges | 818.1456 | JAGURL000000000 |
| jmh44 | N03594 | Blood | CHU Limoges | 818.1484 | JAGWEU000000000 |
| jmh47 | N03736 | Hepatic abscess | CHU Limoges | 818.1421 | JAGURK000000000 |
| jmh50 | Ana-1-p17 | Ascitic fluid | CHU Limoges | 818.1481 | JAGWET000000000 |
| jmh51 | Ana2-p19 | Blood | CHU Limoges | 818.1411 | JAGURJ000000000 |
| jmh58 | Ana4-p37 | Blood | CHU Limoges | 818.1450 | JAGURI000000000 |
| jmh60 | os16-p16 | Hip implant | CHU Limoges | 818.1415 | JAGURH000000000 |
| jmh61 | Ana4-p81 | Continuous ambulatory peritoneal dialysis (CAPD) | CHU Limoges | 818.1480 | JAGWES000000000 |
| jmh62 | 10018 | Peritoneal fluid | CHU Lille | 818.1419 | JAGURG000000000 |
| jmh63 | 10023 | Blood | CHU Lille | 818.1414 | JAGURF000000000 |
| jmh66 | 10254 | Maxilar pus | CHU Lille | 818.1416 | JAGURE000000000 |
| jmh68 | 10263 | Blood | CHU Lille | 818.1451 | JAGURD000000000 |
| jmh71 | 11276 | Blood | CHU Lille | 818.1482 | JAGWER000000000 |
| jmh72 | 11278 | Anal abscess | CHU Lille | 818.1449 | JAGURC000000000 |
| jmh76 | 11285 | Fat tissue | CHU Lille | 818.1483 | JAGYXC000000000 |
| jmh78 | 12309 | Abdominal abscess | CHU Lille | 818.1417 | JAGURB000000000 |
| jmh79 | 11314 | Intestinal cyste | CHU Lille | 818.1447 | JAGURA000000000 |

**Supplementary Table S2.** RNAseq analysis: table of A. all genes, B. upregulated, C. downregulated genes in presence of 0.5% bile, D. COG functional categories enrichment.

**Supplementary Table S3.** BT3563 homologs identified by blastp

| Genome | PATRIC ID | Length (AA) | ALN Length | Identity | Query cover | Subject cover | Hit from | Hit to | Score | E value |
| --- | --- | --- | --- | --- | --- | --- | --- | --- | --- | --- |
| VPI-5482 | fig\|226186.12.peg.3626 (BT 3563) | 293 | 293 | 100 | 100 | 100 | 1 | 293 | 614 | 0 |
| jmh42 | fig\|818.1412.peg.329 | 296 | 296 | 87 | 99 | 99 | 4 | 296 | 532 | 0 |
| jmh43 | fig\|818.1456.peg.161 | 396 | 396 | 34 | 81 | 60 | 149 | 385 | 119 | 8,00E-30 |
| jmh44 | fig\|818.1484.peg.456 | 290 | 290 | 87 | 99 | 100 | 1 | 290 | 525 | 0 |
| jmh47 | fig\|818.1421.peg.2384 | 290 | 290 | 87 | 99 | 100 | 1 | 290 | 525 | 0 |
| jmh50 | fig\|818.1481.peg.2033 | 290 | 290 | 87 | 99 | 100 | 1 | 290 | 525 | 0 |
| jmh51 | fig\|818.1411.peg.850 | 296 | 296 | 87 | 99 | 99 | 4 | 296 | 532 | 0 |
| jmh58 | fig\|818.1450.peg.2100 | 223 | 223 | 97 | 77 | 100 | 1 | 223 | 457 | 1,00E-162 |
| jmh60 | fig\|818.1415.peg.1161 | 399 | 399 | 34 | 81 | 59 | 152 | 388 | 119 | 1,00E-29 |
| jmh61 | fig\|818.1480.peg.1186 | 291 | 291 | 87 | 100 | 100 | 1 | 291 | 527 | 0 |
| jmh62 | fig\|818.1419.peg.360 | 399 | 399 | 34 | 81 | 59 | 152 | 388 | 119 | 9,00E-30 |
| jmh63 | fig\|818.1414.peg.2874 | 396 | 396 | 34 | 81 | 60 | 149 | 385 | 119 | 9,00E-30 |
| jmh66 | fig\|818.1416.peg.3454 | 291 | 291 | 87 | 100 | 100 | 1 | 291 | 527 | 0 |
| jmh68 | fig\|818.1451.peg.696 | 399 | 399 | 34 | 81 | 59 | 152 | 388 | 119 | 1,00E-29 |
| jmh71 | fig\|818.1482.peg.3309 | 399 | 399 | 34 | 81 | 59 | 152 | 388 | 119 | 1,00E-29 |
| jmh72 | fig\|818.1449.peg.2960 | 396 | 396 | 34 | 81 | 60 | 149 | 385 | 119 | 8,00E-30 |
| jmh76 | fig\|818.1483.peg.3838 | 296 | 296 | 85 | 93 | 92 | 26 | 296 | 478 | 1,00E-169 |
| jmh78 | fig\|818.1417.peg.1663 | 293 | 293 | 100 | 100 | 100 | 1 | 293 | 613 | 0 |
| jmh79 | fig\|818.1447.peg.3310 | 293 | 293 | 100 | 100 | 100 | 1 | 293 | 613 | 0 |

**Supplementary Table S4.**
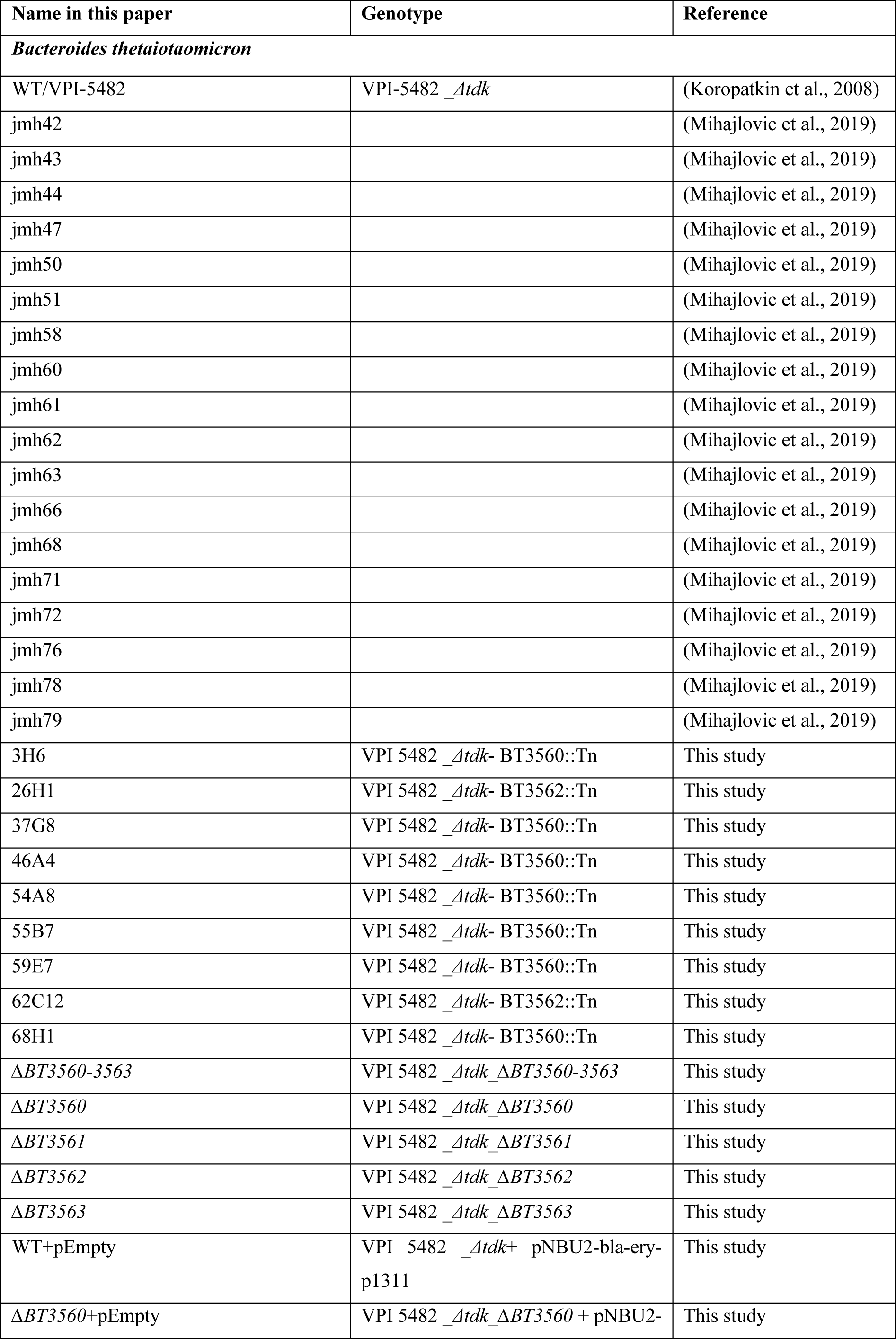
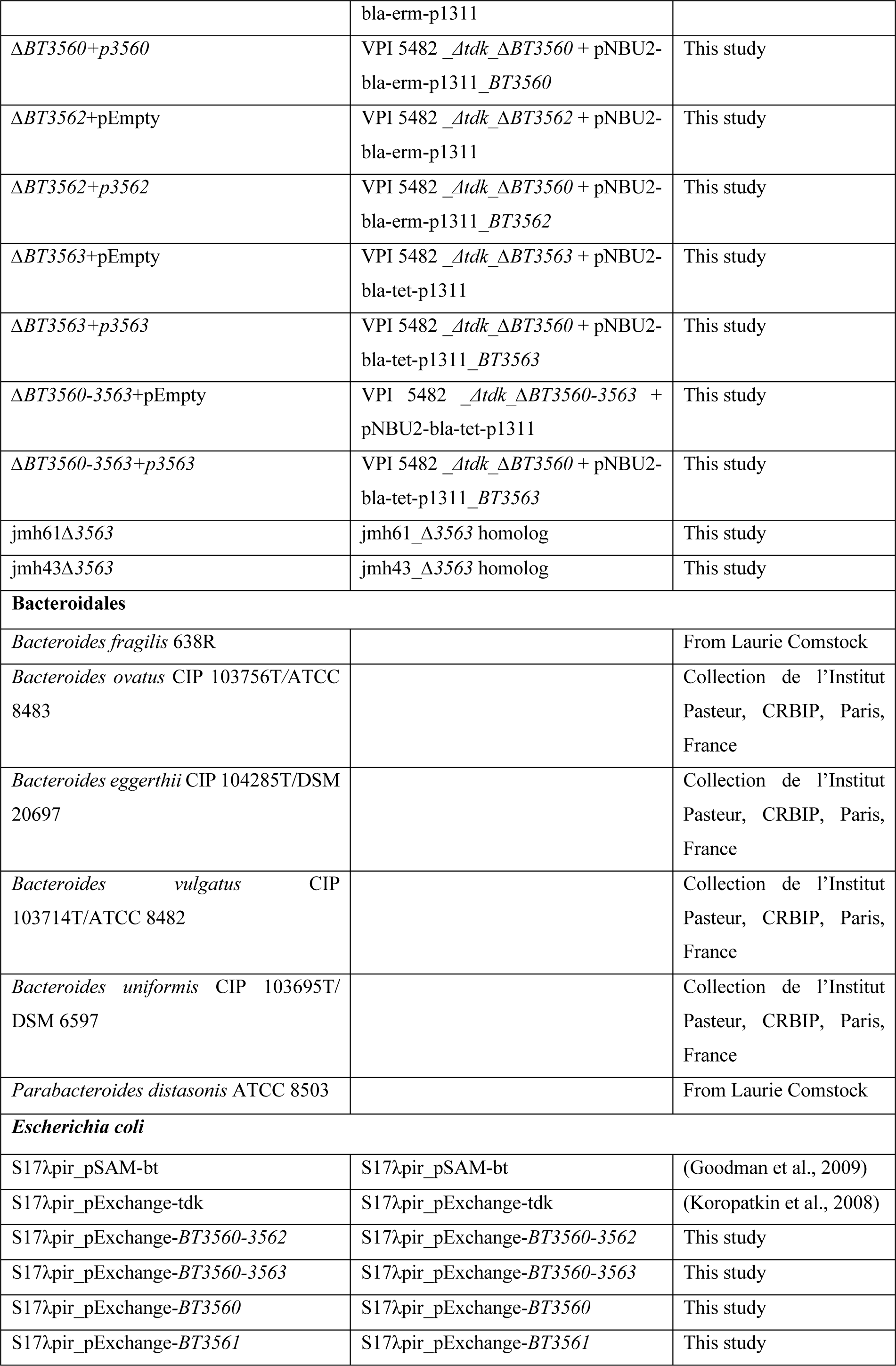
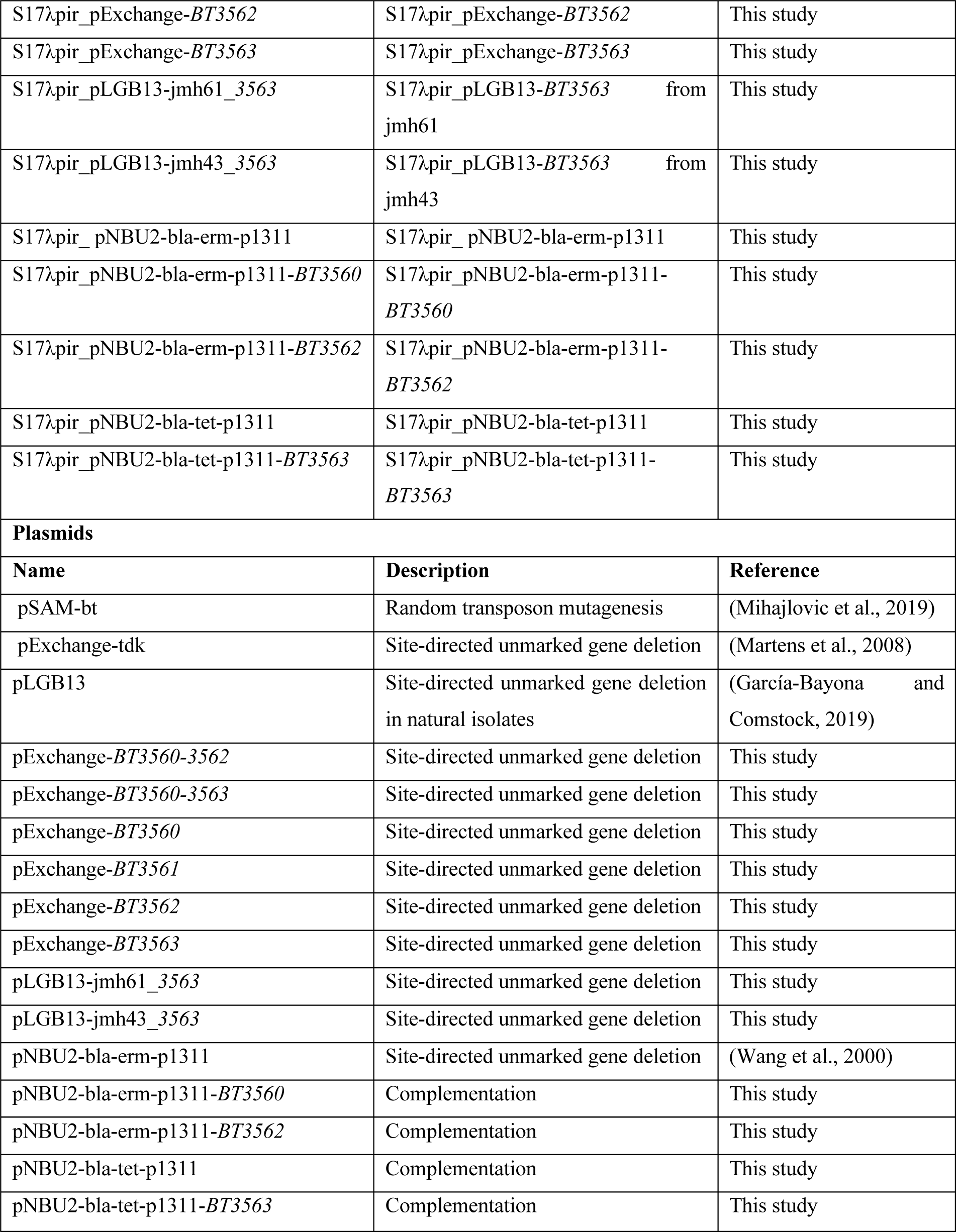
Bacterial strains and plasmids used in this study

**Supplementary Table S5:**
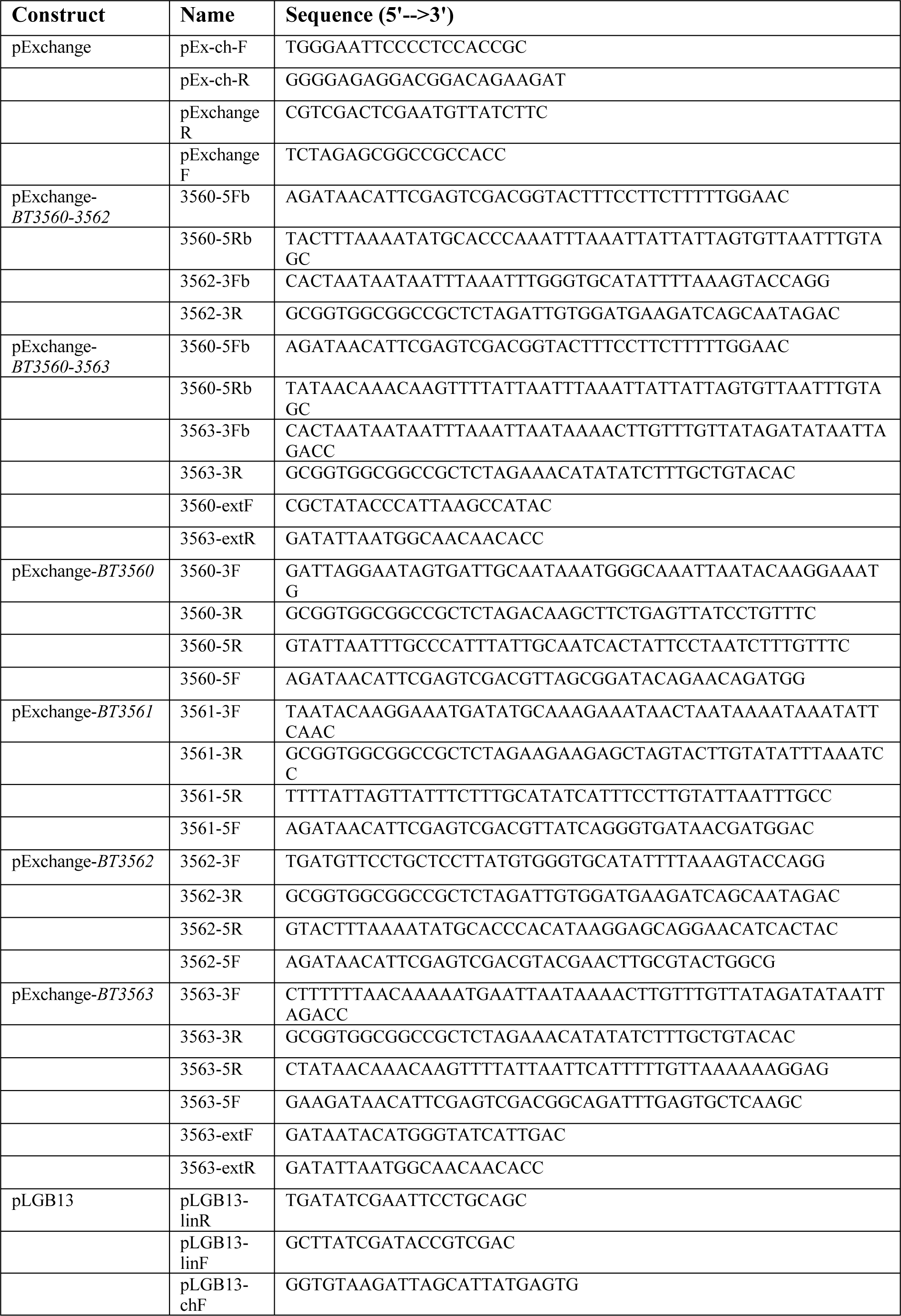
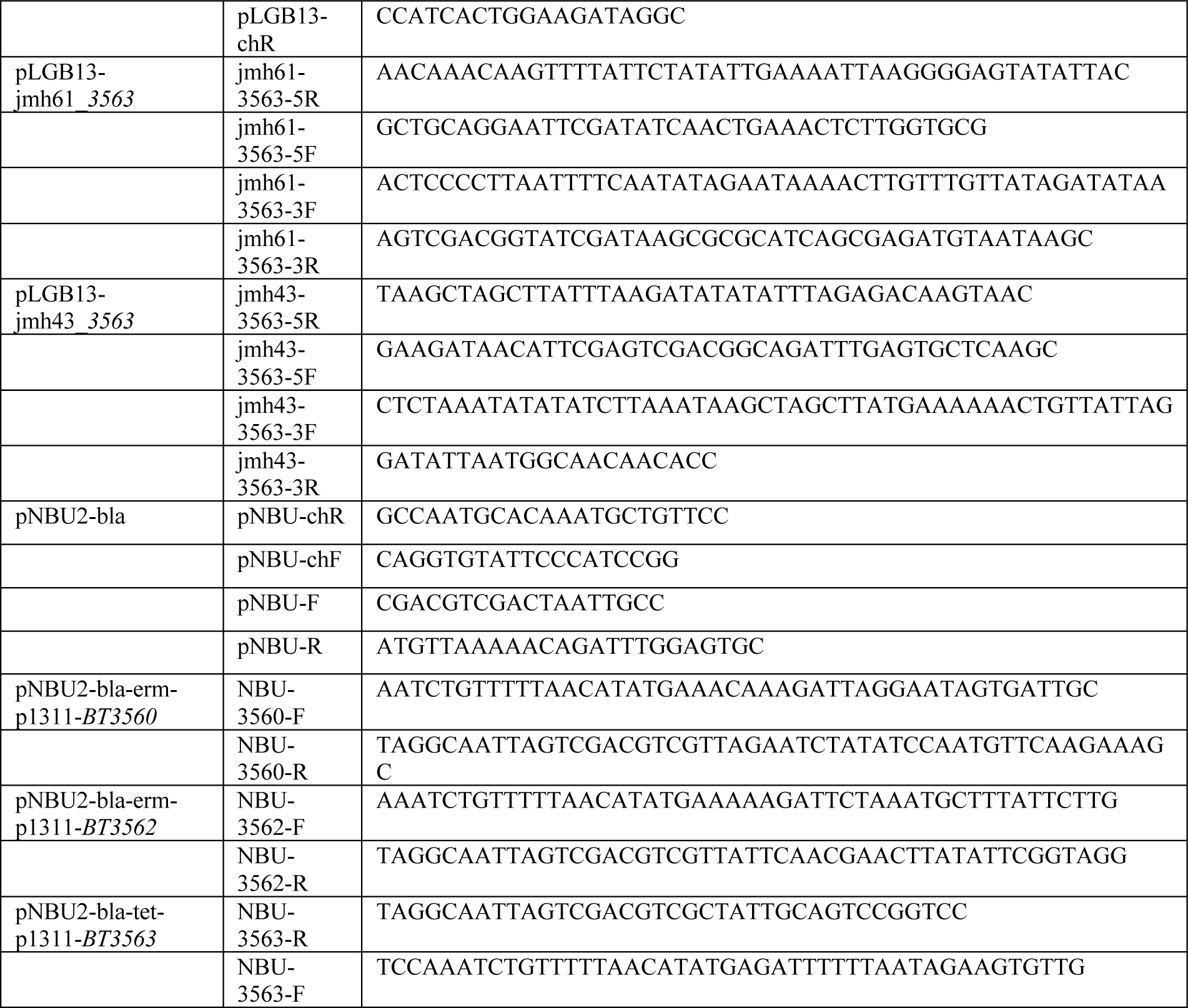
List of primers used in this study.

